# Immortalization of mesenchymal stromal cells by hTERT does not affect the functional properties of secreted extracellular vesicles

**DOI:** 10.1101/2024.12.05.626964

**Authors:** Alessia Brancolini, Madhusudhan Reddy Bobbili, Marianne Pultar, Zahra Mazidi, Matthias Wieser, Johanna Gamauf, Marieke Theodora Roefs, Giulia Corso, Harini Nivarthi, Maria Belen Arteaga Paredes, Dieter Bettelheim, Elsa Arcalis, Sivun Dmitry, Jaroslaw Jacak, Iris Gerner, Matthias Hackl, Johannes Grillari, Regina Grillari-Voglauer

## Abstract

Mesenchymal stromal cell-derived extracellular vesicles (MSC-EVs) have emerged as promising and safe therapeutic agents, however, donor heterogeneities, limited replicative life span and changes in the cellular phenotype throughout in vitro cultivation remain major hurdles for scalable EV production. For these reasons, this study aims to investigate the use of hTERT immortalized (‘telomerized’) MSCs as a potential source for efficient, standardized, reliable MSC-EVs production by comparing parental primary to their telomerized MSC counterparts. We observed that hTERT expression does not affect cell morphology or cellular doubling time, while ensuring unlimited, stable in vitro propagation. In addition, telomerized WJ-MSCs maintained the canonical expression profile of surface markers and the tri-lineage differentiation potential of their primary counterparts. In terms of EV characteristics, the immortalization by hTERT expression did not affect size, number, cargo composition or biological activity regarding anti-inflammatory, anti-fibrotic and wound healing properties in vitro. In summary, the use of hTERT to immortalize MSCs leads to the creation of cell lines that continuously produce MSC-EVs without altering any key functionalities of the cells or resulting EVs. This suggests that telomerization of human cells from single donors is a promising strategy for generating cell factories that can produce EVs in standardized conditions and at scale and with standardization.

## INTRODUCTION

Mesenchymal stromal cells (MSCs) have been proven to have anti-inflammatory, immunomodulatory and regenerative capabilities in various pre-clinical models of immunological and degenerative diseases (Galipeau and Sensébé, 2018; Squillaro et al., 2016; Trento et al., 2018). While it is now evident that the activities of MSCs in vivo are primarily mediated by paracrine factors, many, if not the majority, of the regenerative effects observed might be mediated by the release of extracellular vesicles (EVs) (Deng et al., 2015; Wang et al., 2015). Indeed, EVs obtained from mesenchymal stromal cells (MSC-EVs) isolated from various donors and tissues, with different culture conditions and enrichment strategies, have proven to be effective in pre-clinical models, exhibiting therapeutic benefits similar to those attributed to the parent cells. Moreover, MSC-EVs offer several advantages over cell-based therapies, reducing some technical and safety issues (Harrell et al., 2019; Lener et al., 2015; Lotfy et al., 2023; Lou et al., 2017; Witwer et al., 2019; Yaghoubi et al., 2019). For example, MSC-EVs exclude the possibility of tumor formation since they lack the ability to self-replicate (Volarevic et al., 2018). The likelihood of immunological reactions is also decreased by their small size and limited expression of membrane histocompatibility complexes (Lai et al., 2019; Reis et al., 2016). Furthermore, MSC-EVs can enter the circulation, quickly migrate into different tissues (Shelke et al., 2014) and cross the blood-brain barrier (BBB) (Xin et al., 2013), whereas MSCs retain mainly in the lung (Wang et al., 2010) and have difficulty crossing the BBB (Cerri et al., 2015). For these reasons, MSC-EVs are currently considered suitable candidates for a variety of therapeutic purposes, as demonstrated by the number of pre-clinical and clinical trials currently ongoing (Birtwistle et al., 2021; Cosenza et al., 2018; Doeppner et al., 2015; Long et al., 2017; Lotfy et al., 2023; Nojehdehi et al., 2018; Xue et al., 2021; Yaghoubi et al., 2019).

Such large potential therapeutic need of single EV-doses, however, clearly shows that scalable and standardized EV production is a pre-requisite for clinical translation. This is currently hampered by the limited replicative life span of primary MSCs, which leads to the necessity of donor pooling for securing sufficient cell numbers for generating master cell banks for biotechnological production processes. Moreover, changes in the cellular phenotype throughout in vitro cultivation generate further challenges when using human primary cells (Burnouf et al., 2019; Lener et al., 2015; Maumus et al., 2020; Rayyan et al., 2018). Similarly, EVs from MSCs isolated from different donors have been found to exhibit profound cargo and functional variability (Giebel, 2017; Madel et al., 2023; McBride et al., 2021; Ragni et al., 2019; Scheiber et al., 2022). The functional changes of MSC-EVs caused by differences in their cargo content raise the issue of standardization and reproducibility when using EV-based therapies (Kordelas et al., 2019, 2014). Moreover, a key step in the transition from research to the development of clinically approved EV-based products is the scalability of EVs (Gowen et al., 2020). Several strategies have been developed for this purpose either based on physical or chemical stimulation of the cells to increase particle secretion (Wang et al., 2020; Zhu et al., 2018) or by exploiting large-scale cell culture platforms. Indeed, culturing MSCs in large-scale 3D bioreactors, has been demonstrated to produce noticeably higher yields of EVs compared to two-dimensional cell cultures (Gobin et al., 2021; Haraszti et al., 2018; Tacheny, 2021).

However, regardless of the EV production strategy exploited, the first step of scalable EV manufacturing is adequate cell expansion, which is hindered by the limited proliferative potential of adult tissue-derived MSCs. On the one hand, long-term serial passaging of cells to reach necessary quantities of EVs may lead to loss of clonal composition and differentiation capacity of the culture, thus affecting EVs potency (McKee and Chaudhry, 2017). On the other hand, replicative senescence after prolonged expansion may decrease the therapeutic efficacy of EVs secreted by high-passage cells nearing replicative senescence (Almeria et al., 2022; Dorronsoro et al., 2021; Robbins, 2017) or worse, may even have detrimental effects (Dorronsoro et al., 2021). Indeed, it has been proven that EVs from senescent MSCs have decreased osteogenic potential compared to EVs from early passage MSCs, and their cargo composition reflects the senescence state of MSCs (Lei et al., 2017).

To circumvent replicative senescence, various strategies have been employed over the last decades, such as using cells isolated from tumors (Soule et al., 1990), waiting for spontaneous immortalizing mutations in primary cell cultures (Harvey and Levine, 1991; Lehman et al., 1993), shortening the wait using mutagens (Busuttil et al., 2003; Caliri et al., 2020; Crane, 1999), viral or cellular oncogenes or combinations thereof (Chalak et al., 2024; Linder and Marshall, 1990). While all these strategies have led to successful immortalizations, they come at the expense of losing primary cell characteristics. In contrast, Bodnar et al were the first to demonstrate that by transfecting human retinal pigment epithelial cells and foreskin fibroblasts with the catalytic subunit of human telomerase (hTERT), which assembles into the holoenzyme automatically together with the constitutively transcribed RNA component of telomerase (hTERC), it is possible to prevent telomere shortening and thus immortalize primary human cells (Bodnar et al., 1998). Therefore, Shay and Wright have suggested to use the term ‘telomerized cellś and foresaw their use in biotechnological applications (Shay and Wright, 2007). By now, a plethora of studies have shown that the primary characteristics of many different cell types after immortalization by hTERT overexpression remain highly similar to their parental primary cells (Bachand et al., 2000; Chang et al., 2005; Kassem et al., 2004; Kharbanda et al., 2000; Simonsen et al., 2002; Twine et al., 2018; Wagner et al., 2018; Wieser et al., 2008; Wolbank et al., 2009), not affecting cellular karyotype, differentiation properties, contact-inhibition nor do they display a tumorigenic phenotype for up to 100 population doublings in vitro (Chang et al., 2005; Kitagawa et al., 2007; Matsumura et al., 2004; Wieser et al., 2008). Telomerized endothelial cells even integrated well into mouse vasculature (Yang et al., 2001), while the therapeutic potential of telomerized MSCs has been shown both in vitro (Kobune et al., 2005), and in vivo in different mouse models (Hornsby, 2004; Yang et al., 2001).

Still, so far, the effects of hTERT immortalization of parental cells on EV properties have not been studied. Here, we compared primary versus telomerized Wharton’s Jelly-derived MSCs (WJ-MSC) along with their secreted EVs to assess several key biomarkers and functionalities in vitro. Our findings suggest that using telomerized MSCs as a cell factory for EV production is a viable and efficient approach for producing standardized and scalable amounts of MSC-EVs, which is currently in high demand (Ng et al., 2022; Whitford and Guterstam, 2019).

## MATERIALS AND METHODS

### 2.1 Cells and Culture Conditions

#### 2.1.1. Cell line establishment

Human mesenchymal stromal cells were isolated from Whartońs Jelly with ethical approval from the responsible Institutional Review Board (Medical University of Vienna) and written informed consent by the donor aged 32 with no clinical history. Cells were isolated following published protocols (Beeravolu et al., 2017) within 24 hours after surgery using mechanical dissociation. Thereafter, cells were seeded into culture flasks pre-coated with Animal component-free cell attachment substrate (ACF, Stemcell Technologies) in Mesencult^TM^ ACF Plus (Stemcell Technologies) with 2 mM GlutaMAX-I^TM^ (Gibco) and Mesencult^TM^ ACF Plus 500X supplements (Stemcell Technologies) and incubated at 37°C in a humidified atmosphere with 5% CO_2_. After 21 culturing days and upon having reached about 70% confluence, the primary cells WJ-MSC273 were transfected by electroporation with a plasmid carrying hTERT-coding cDNA, which was excised from plasmid pGRN145 (kindly provided by Geron, Menlo Park, CA), and neomycin-phosphotransferase. Positively transfected cells were selected using 200 µg/mL Geneticin sulfate (G418, InvivoGen) and enriched for the establishment of a master cell bank (WJ-MSC/TERT273).

##### Cell Lines Maintenance

WJ-MSC/TERT273 cells were passaged when having reached about 80-90% confluence. Therefore, the cells were washed twice with PBS and detached from the culture flasks by incubation with CTS^TM^ TrypLE^TM^ Select Enzyme solution (20 µL/cm^2^, Gibco) at 37°C until detachment. Thereafter, the cell suspension was harvested and centrifuged for 5 minutes at 300 x g. Cells were seeded into new ACF pre-treated flasks with a 1:3 to 1:6 split ratio and cultivated at 37°C in a humidified atmosphere with 5% CO_2_. HEK293 cells were propagated in MEM Eagle, w: 1.5 g/L NaHCO_3_ (PAN Biotech) supplemented with 2 mM L-glutamine (Gibco), 10% fetal bovine serum (FBS, Sigma), 1 mM Na-pyruvate, and nonessential amino acids (Gibco) at a splitting ratio of 1:8 to 1:12 twice a week following the manufacturer’s instruction. RAW264.7 mouse macrophages (ATCC) were propagated in Minimum Essential Medium Eagle (Sigma) supplemented with 1X GlutaMAX^TM^-I (Gibco) and 10% FBS (PAN Biotech) at a splitting ratio of 1:9 to 1:18 twice a week following the manufacturer’s instruction. Human foreskin fibroblasts fHDF/TERT166 (Evercyte GmbH) were propagated in DMEM-Ham’s F12 (1:1), w: stable Glutamine, w: 1.2 g/L NaHCO_3_ (PAN Biotech) supplemented with 10 % FBS (PAN Biotech) and 100 μg/mL G418 (InvivoGen). Cells were passaged twice a week with a split ratio of 1:4 following the protocols provided by the manufacturer. Telomerized human umbilical vein endothelial cells HUVEC/TERT2 (Evercyte GmbH) were propagated in EBM^TM^ (Lonza) supplemented with BBE, hEGF, Hydrocortisone, Ascorbic acid (components of EGM^TM^ SingleQuots^TM^ supplement kit, Lonza) 10% FBS (PAN Biotech) and 20 µg/mL G418 (InvivoGen). Cells were propagated with a split ratio of 1:8 twice a week on Gelatine (Sigma) pre-coated flasks following the protocols provided by Evercyte.

### 2.2 Cell characterization

#### 2.2.1 Analysis of presence of typical surface marker proteins

The expression of the typical surface marker proteins on WJ-MSC/TERT273 (telomerized) and WJ-MSC273 (primary) was assessed by performing immunofluorescent staining and a flow cytometric analysis. Therefore, the following conjugated monoclonal antibodies from BD Pharmingen were used: APC mouse anti-human CD34 (1:10 dilution in blocking solution), APC mouse anti-human CD73 (1:40 dilution in blocking solution), APC mouse anti-human CD90 (1:80 dilution in blocking solution), APC mouse anti-human CD105 (1:40 dilution in blocking solution). 3 × 10^5^ cells were resuspended in PBS, incubated with 200 µL/sample in blocking solution (10% FBS in PBS) for 15 minutes at 37°C, followed by incubation with the monoclonal antibodies for 30 minutes at 37°C protected from light. After a washing step with blocking solution cells were resuspended in 400 µL of DAPI solution (Roche) (1:50 dilution in PBS). As a negative control, cells were incubated with the isotype control antibody APC mouse IgG1 (BD Pharmingen, 0.2 mg/mL). Then, the cells were analyzed using the ZE5 Cell analyzer (Bio-Rad) by setting a gate at 10,000 events for viable cells. Data analysis was performed using Kaluza Analysis 2.1 software.

#### 2.2.2 Multilineage Differentiation of MSCs

To assess the osteogenic differentiation ability of WJ-MSC/TERT273 (telomerized) and WJ-MSC273 (primary), 1.8 x 10^4^ cells/well were seeded into a pre-coated 24-well plate (see 2.1.2). After 24 hours, the medium was changed to either osteogenic differentiation medium or control medium, followed by a medium change twice a week until staining. Osteogenic differentiation medium was prepared by supplementing DMEM low glucose medium (Gibco) with 10% FBS (PAN Biotech), L-ascorbic Acid 2-phosphante sesquimagnesium salt hydrate 150 µM (Sigma), B1a25-Dihydroxyvitamin D3 1uM (Sigma), Glycerophosphate disodium salt hydrate 10 mM (Sigma), Primocin 100 ug/mL (InvivoGen) and Dexamethasone 1 µM (Sigma), whereby Dexamethasone was always added freshly to the medium. The control medium was prepared by adding 10% FBS (PAN Biotech) and L-Glutamine 1.78 mM (Gibco) to DMEM low glucose (Gibco)/Ham’s F12 (Gibco) 1:1 mixture. After 15 days, matrix mineralization was measured by cell fixation using 70% Ethanol at -20°C for 30 minutes, followed by incubation with Alizarin red staining solution (AR, 14 mg/mL of Alizarin red S (Sigma) dissolved in HQ Water at 30°C for 30 minutes) for 10 minutes at RT. Thereafter, cells were incubated with 200 µL/well of 0.5% SDS / 0.1 M HCl-HQ water solution for 30 minutes at RT. Quantification of matrix mineralization was performed by transferring 50 µL of the resulting solution into a 96 well plate and measuring at 425 nm wavelength. Measurements were carried out in triplicates for each well and in quintuplicates for each biological sample analyzed. To assess the adipogenic differentiation ability of WJ-MSC/TERT273 (telomerized) and WJ-MSC273 (primary) cells, 1.4 x 10^4^ cells/well were seeded into a pre-coated 24-well plate (see 2.1.2). A medium change to either adipogenic differentiation medium or control medium was performed 24 hours after seeding, which then was repeated twice a week until staining. Adipogenic differentiation medium was prepared by supplementing DMEM high glucose (Gibco) / Ham’s F12 (Gibco) 1:1 mixture with 10% FBS (PAN Biotech), Indomethacin 100 µM (Sigma), Insulin 10 µg/mL (Sigma), IBMX 50 µM (Sigma), Primocin 100 µg/mL (InvivoGen) and Dexamethasone 1µM (Sigma), which was added freshly before treatment. The control medium was prepared by adding 10% FBS (PAN Biotech) and L-glutamine 1.78 mM (Gibco) to DMEM high glucose (Gibco)/Ham’s F12 (Gibco) 1:1 mixture. For staining developed lipid droplets at day 21, Oil Red O Solution was prepared by diluting 6 mL of Oil Red O stock solution (Sigma) with 4 mL HQ Water, followed by incubation at RT for 15 minutes and passing through a 0.45 µm filter. Cells were stained following fixation with Roti-Histofix (4% paraformaldehyde, Carl Roth) at RT for 10 minutes. Thereafter, the cells were incubated with 100% isopropanol for 10 minutes to dissolve the stained lipid droplets. Quantification of droplet formation was performed by transferring 150 µL of solution into a 96 well plate and measuring at 520 nm wavelength. Measurements were carried out in triplicates for each well and in quintuplicates for each biological sample analyzed. The spectrophotometric quantification of both Alizarin Red and Oil Red O staining was performed using a Spark^®^ multimode microplate reader (Tecan), images were taken using EVOS™ XL Core microscope (EVOS). Chondrogenic differentiation was performed by pelleting 200.000 cells at 330 x g in a 15 mL falcon tube. The Gibco StemPro^TM^ Chondrogenesis Differentiation Kit (Thermo Fisher Scientific), consisting of the StemPro^TM^ Osteocyte/Chondrocyte Differentiation Basal Medium and the StemPro^TM^ Chondrogenesis Supplements, was used for induction of differentiation as described by the manufacturer. In brief, 1% PenStrep (Szabo Scandic) was added to a mixture of 90 mL of basal medium and 10 mL of supplements. Differentiation medium (1 mL) was used as solution in which the cells were pelleted. The medium was changed twice a week. To avoid the pellets adhering to the falcon tube surface, tubes were carefully agitated each day. After 21 days, the pellets were rinsed twice with PBS without Ca^++^/MG^++^ and fixed with 4% paraformaldehyde (Herba Chemosan). After 48 hours, 70% alcohol was used to replace the paraformaldehyde, and the pellets were then embedded in paraffin and sectioned. To verify the synthesis of acidic mucosubstances, sections were stained with Alcian blue (Waldeck GmbH) in accordance with a standard staining methodology.

#### 2.2.3 Measurement of telomerase activity

For determination of telomerase activity (TA), a modified real-time telomeric repeat amplification protocol (TRAPeze RT Telomerase Detection Kit, Millipore) was used and TA was calculated relative to internal Control Template TSR8 (0.2 amols/µL). Cell extracts were prepared by detaching WJ-MSC273 (primary) and WJ-MSC/TERT273 (telomerized) cells from the culture flask according to cell maintenance protocols (2.1.2); cells were counted and pelleted at 200.000 cells/condition. Pellets were washed twice with PBS, re-pelleted and immediately lysed in CHAPS Lysis Buffer (supplied by the kit). The cell lysate was incubated on ice for 30 minutes and spun at 12.000 x g for 20 minutes at 4°C. The supernatant was then used for PCR Amplification according to the protocol specifications. Telomerase activity was further assessed by TeloTAGGG Telomerase PCR ELISA PLUS assay (Roche), and TA was calculated relative to internal control template TSR8 (1 amols/µL). Cell extracts were prepared as described above, and 200 µL of lysis reagent (supplied by the kit) was used for 2 x 10^5^ cells (1000 cells/µL lysis reagent). The supernatant was then used for Telomeric repeat amplification (TRAP reaction) according to the manufacturer’s instructions.

#### 2.2.4 Staining for SA-β-Galactosidase activity

2.5 x 10^4^ cells/well were seeded into a pre-coated 24-well plate (see 2.1.2). After 24 hours, the supernatant was removed, cells were washed twice with PBS and fixed for 10 minutes at RT with 0.5 mL/well of Fixative solution prepared by mixing 2.5 mL of PBS, 2.5 mL of Roti-Histofix (4% paraformaldehyde, Carl Roth) and 40 µL of 25% Glutaraldehyde (Sigma). Cells were washed twice with PBS and once with 500 µL/well of X-Gal washing buffer prepared by mixing 2.5 mL of 504 mM Na_2_HPO_4_ (Sigma) 2.5 mL of 148 mM Citric Acid (Sigma) and 5.0 mL of HQ water. The staining buffer was prepared by mixing 50 µL 400 mM potassium hexacyanido-ferrate (II) (Sigma), 50 µL of 400 mM potassium hexacyno-ferrate (III) (Sigma), 40 µL of 200 mM MgCl2, 200 µL of 20 mg/mL X-Gal (Roche) dissolved in DMF (Sigma) and 3660 µL of X-Gal washing buffer. The plate was sealed with parafilm and incubated at 37°C / 5% CO_2_ for 16 hours. Evaluation of SA-ß-Galactosidase activity was done microscopically and images were taken using EVOS™ XL Core microscope (EVOS). The percentage of positive cells was calculated by manual count of stained cells over the total number of cells per microscopy field.

#### 2.2.5 Evaluation of Tumorigenic Potential by Soft Agar Assay

The tumorigenic potential of WJ-MSC273 (primary) and WJ-MSC/TERT273 (telomerized) cells was evaluated using CytoSelect™ 96-Well Cell Transformation Assay (Soft Agar Colony Formation, Cell Biolabs). The preparation of the base and cell agar layers was performed according to the manufacturer’s instruction, and 2× cell culture medium (Mesencult^TM^ ACF Plus (Stemcell Technologies) with 4 mM GlutaMAX-I^TM^ (Gibco) and Mesencult^TM^ ACF Plus 2500X supplement (Stemcell Technologies) was used for agar layer preparation. To set the optimal cell number, the cells were counted using a hemocytometer and adjusted to 1.2 x 10^5^ and 3.6 x 10^5^ cells/mL in complete medium (the assay was performed in triplicates). Then, the plates were incubated in a humified incubator for 6 days and analyzed for colony formation. The quantitation of anchorage-independent growth was performed according to the manufacturer’s instruction by cell solubilization, lysis and detection by CyQuant^®^ GR Dye. The spectrophotometric quantification of Anchorage-Independent Growth was performed using a Spark^®^ multimode microplate reader (Tecan), images were taken using EVOS™ XL Core microscope (EVOS).

### 2.3 EV preparation and molecular characterization

#### 2.3.1 Extracellular vesicle enrichment

For EV production, WJ-MSC273 (primary) and WJ-MSC/TERT273 (telomerized) cells were cultured for 78 hours in G418-free Mesencult^TM^ ACF Plus medium. When cells reached a confluency of 75–85%, supernatant was collected, and cell density and viability were determined with Trypan blue exclusion method using Cell-Chip Hemocytometer (Bioswisstec). The conditioned medium was centrifuged at 700 x g for 5 minutes at RT. The supernatant was further centrifuged at 2000 x g for 10 minutes at RT and filtered through a 0.22-μm PVDF-filter to exclude larger particles. The supernatant was stored in aliquots at -80°C for up to 4 weeks before EV enrichment. Hollow fiber columns with 300 kDa cut-off (MicroKros 20CM 300K MPES 0.5MM MLL X FLL 1/PK STER, Repligen) were used for TFF enrichment of EVs. Sample concentration was alternated with ice-cold HEPES washes (Sigma, 20 mM in cell grade H_2_O and 0.22 µm filtered) throughout the process and the sample (from 50 to 100 mL starting volume) was concentrated down to 1 mL and stored in aliquots at -80°C until further analysis. For the medium control used in the biological assays, the same volume of unconditioned Mesencult^TM^ ACF Plus medium was processed following the protocol described above.

#### 2.3.2 Analysis of particle number and size distribution

Nanoparticle tracking analysis (NTA) was used to evaluate the concentration, size distribution, median and mode sizes of EVs using the ZetaView BASIC PMX-120 (Particle Metrix GmbH). Particle count and size distribution were calculated using ZetaView software (version 8.05). The ZetaView device was calibrated using the provided standard beads; for the analysis of the EV preparations, the sensitivity was set at 80, the shutter was set at 100, temperature was set at 23.0 °C and the frame rate was set at 30. The EV samples were diluted with filtered PBS prior to particle measurement. For each particle measurement, three replicates were recorded and analyzed.

#### 2.3.3 Transmission electron microscopy (TEM)

To visualize primary and TERT EVs via electron microscopy, samples were prepared as described previously (Hausjell et al., 2023). Formvar-coated copper grids were floated on a 10 μL sample drop (0.6-1.1 x 10^9^ particles) for 5 minutes. After gently removing the excess of sample with filter paper (Whatman), grids were quickly washed with distilled water and subsequently floated in a drop of 2% glutaraldehyde for 10 minutes for fixation. Following a quick wash with distilled water, negative staining was performed with uranyl acetate replacement stain (UAR_EMS Stain, EMS-Electron Microscopy Sciences) by floating the grids on a drop of UAR twice for 10 seconds and a third time for 60 seconds, gently blotting off the excess of stain after each step. Grids were let air dry prior to imaging using a FEI Tecnai G2 transmission electron microscope operating at 160 kV.

#### 2.3.4 Protein content quantification

Bicinchoninic Acid (BCA) protein assay (Thermo Fisher Scientific) was used to quantify the protein concentration in the WJ-MSC273 and WJ-MSC/TERT273-derived EV preparations according to the manufacturer’s instructions. EVs were diluted (1:25 to 1:50) in filtered PBS to a final EV concentration of 5 x 10^9^ particles/mL prior to protein measurement. Using a Spark® multimode microplate reader (Tecan), absorbance was measured at 562 nm. The purity of EV preparations was assessed by calculating the ratio between particle number, as indicated by NTA, and protein content, as evaluated by BCA assay.

#### 2.3.5 Analysis of presence of EV marker proteins by flow cytometric analysis

The analysis of classical tetraspanin markers on WJ-MSC273 and WJ-MSC/TERT273-derived EVs was carried out using exosome-human CD81 beads (Invitrogen) to capture EVs, followed by antibody staining and flow cytometric analysis. Briefly, EV samples were diluted to a concentration of 1 x 10^9^ particles in 100 µL of filtered PBS and incubated with the CD81+ beads (1:6 dilution) overnight at +4°C on a plate shaker (500 rpm). The bead-EV complexes were washed with 700 µL/sample of filtered PBS and collected after incubation for 5 minutes at RT in a magnetic separator by carefully removing the supernatant. The bead-EV complexes were resuspended in 100 µL/sample of filtered PBS and incubated with the following conjugated monoclonal antibodies from Miltenyi Biotec: mouse anti-CD9-APC antibody (Clone REA1071), anti-CD63-FITC antibody (Clone REA1055) and anti-CD81-APC (Clone REA513). 1 μL of antibody was added to each sample (1:100 dilution) and samples were incubated at 4°C for 1 hour on a shaker (500 rpm). Then, 700 µL of filtered PBS were added to each sample and the EV-bead complexes were collected by incubation for 5 minutes at RT in a magnetic separator; supernatant was removed, and pellets were resuspended in 200 μL of filtered PBS. The samples were analyzed using the ZE5 Cell analyzer (Bio-Rad), with a gate set at 10,000 events for acquisition. Data analysis was performed using Kaluza Analysis 2.1 software.

#### 2.3.6 EV surface protein profiling by Multiplex bead-based flow cytometry assay

Surface protein profiling of primary and TERT EVs was performed via Multiplex bead-based EV analysis followed by a flow cytometric analysis (MACSPlex EV IO Kit, human, 130-108-813, Miltenyi Biotec). EV-containing samples were processed as follows: 2.5 x 10^9^ particles were diluted with MACSPlex buffer (MPB) to a final volume of 120 µL and loaded onto wells of a pre-wet and drained MACSPlex 96-well 0.22 µm filter plate. 15 µL of MACSPlex Exosome Capture Beads (containing 39 different antibody-coated bead subsets) were added to each well. Filter plates were then incubated on an orbital shaker overnight (14–16 hours) at 450 rpm at RT protected from light. To wash the beads, 200 µL of MPB was added to each well and centrifuge 300 x g at RT for 3 min. For counterstaining of EVs bound by capture beads with detection antibodies, 135 µL of MPB and 5 µL of each APC-conjugated detection antibody cocktail (anti-CD9, anti-CD63, and anti-CD81) were added to each well and plates were incubated on an orbital shaker at 450 rpm protected from light for 1 hour at RT. Next, plates were washed twice by adding 200 µL MPB to each well and centrifuge 300 x g at RT for 3 minu. Add 200 µL MPB to each well and incubate on an orbital shaker at 450 rpm protected from light for 15 minutes at RT. Subsequently, wells were drained and 200 µL of MPB was added to each well. Beads were resuspended by pipetting and transferred to V-bottom 96-well microtiter plate (Thermo Fisher Scientific). Flow cytometric analysis was performed using CytoFLEX S (Beckman Coulter). All samples were automatically mixed and 150–170 µL of samples were immediately loaded to and acquired by the instrument, resulting in approximately 3,000–5,000 single bead events recorded per well. CytExpert software (Beckman Coulter) was used to analyze flow cytometric data. Median fluorescence intensity (MFI) for all 39 capture bead subsets were background corrected by subtracting respective MFI values from matched non-EV buffer (PBS) that were treated as EV-containing samples (buffer/medium + capture beads + antibodies).

#### 2.3.7 Analysis of EV marker purity by Western Blot analysis

WJ-MSC273 and WJ-MSC/TERT273 cells were collected, and the cell pellets (1.5 x 10^6^ cells) were lysed with 150 µL of 1× RIPA buffer (Merck Millipore), kept on ice, and vortexed five times every 10 minutes for 2 hours. The cell lysate was then centrifuged at 15,000 × g for 5 minutes at 4°C and the supernatant was collected. Protein concentrations in the supernatants were quantified by BCA assay (Thermo Fisher Scientific) according to the manufacturer’s instructions. 1 x 10^10^ particles from WJ-MSC273 and WJ-MSC/ TERT273 derived EVs were diluted in 0.22 µm filtered PBS to a final volume of 200 µL. 25 µL of Exosome-Human CD81/CD9/CD63 beads (Invitrogen), were added to each EV preparation. Primary and TERT EVs were incubated for beads capture on a shaker set on 500 rpm at 4°C overnight. Incubation was followed by a wash with filtered PBS and incubation on a DynaMagTM magentic rack (Thermo Fisher Scientific). Beads were resuspended in 30 µL of filtered PBS. 10 µg of cell lysates and the recovered EV particles were mixed with NuPAGE™ LDS ample Buffer (Thermo Fisher Scientific) and NuPAGE™ Sample Reducing Agent (Thermo Fisher Scientific) and heated at 90°C for 10 minutes. Proteins were separated on a NuPAGE Novex 4–12% Bis-Tris Protein Gel (Thermo Fisher Scientific) in NuPAGE™ MOPS SDS Running Buffer (Thermo Fisher Scientific) along with a Chameleon® Duo Pre-stained Protein Ladder (Licor). The proteins on the gel were transferred to a Trans-Blot Turbo Midi 0.2 µm PVDF Transfer Packs (Bio-Rad) nitrocellulose membrane using the Mix MW program of the Trans-Blot Turbo Transfer System. The membrane was stained with Ponceau S solution (Sigma Aldrich) and blocked with 50% Intercept® (TBS) Blocking Buffer (Licor) in TBS (Thermo Fisher Scientific). After blocking, the membrane was incubated overnight at 4°C with primary antibodies in v/v 50%/50% Intercept® (TBS) Blocking Buffer (Licor)/TBS (Thermo Fisher Scientific) supplemented with 0.1% Tween-20 (Sigma Aldrich). The following primary antibodies were used: TSG101, abcam, ab125011, 1:1,000; Syntenin-1, Origene, TA504796, 1:1,000; Calnexin, GeneTex, GTX101676, 1:1000, CD81 (B-11), Santa Cruz, sc-16602. The membrane was washed three times with TBS (Thermo Fisher Scientific) supplemented with 0.1% of Tween-20 (TBS-T) and incubated with the corresponding secondary antibody (LI-COR; anti-mouse IgG, 926-68072, 1:7500; anti-rabbit IgG, 925-32213, 1:7500) for 1 hour at RT. Finally, the membrane was washed with TBS-T and visualized with the BioRad ChemiDoc imaging system at 680 and 800 nm.

#### 2.3.8 MiRNome Profiling

1 x 10^9^ particles were used for total RNA extraction from WJ-MSC273 and WJ-MSC/ TERT273 derived EVs. Small RNA-seq libraries were generated using the miND® workflow (Khamina et al., 2022). Therefore, 8.5 μL total RNA were mixed with a set of 7 spike-in oligonucleotides harboring a unique 13mer core and 4N-randomized ends to enable extensive quality control and absolute normalization. Libraries were generated using the RealSeq Biofluids kit, pooled, and sequenced on an Illumina NextSeq550 (75bp, single-end). NGS data were analyzed using the miND® pipeline (TAmiRNA) (Diendorfer et al., 2022). The overall quality of the next-generation sequencing data was evaluated automatically and manually with fastQC v0.11.9 (Andrews 2010) and multiQC v1.10 (Ewels et al. 2016). Reads from all passing samples were adapter trimmed and quality filtered using cutadapt v3.3 (Martin 2011) and filtered for a minimum length of 17nt. Mapping steps were performed with bowtie v1.3.0 (Langmead et al. 2009) and miRDeep2 v2.0.1.2 (Friedländer et al. 2012), whereas reads were mapped first against the genomic reference GRCh38.p12 provided by Ensembl (Zerbino et al., 2018) allowing for two mismatches and subsequently miRBase v22.1 (Griffiths-Jones, 2004), filtered for miRNAs of hsa only, allowing for one mismatch. For a general RNA composition overview, non-miRNA mapped reads were mapped against RNAcentral (The RNAcentral Consortium, 2019) and then assigned to various RNA species of interest. Statistical analysis of preprocessed NGS data was done with R v4.0. Differential expression analysis with edgeR v3.32 (Robinson et al., 2010) used the quasi-likelihood negative binomial generalized log-linear model functions provided by the package. The independent filtering method of DESeq2 (Love et al., 2014) was adapted for use with edgeR to remove low abundante miRNAs and thus optimize the false discovery rate (FDR) correction. Correlation analysis is based on rpm normalized miRNA counts. The mean rpm over the three replicates was calculated and Spearman rank test was performed. To check for telomerase specific sequences in the small RNA sequencing data, trimmed and filtered reads were mapped against gene and transcript sequences (NG_009265.1, NM_001193376.3 and NM_198253.3) using bowtie v1.3.0 (Langmead et al., 2009).

#### 2.3.9 Genomic DNA isolation and hTERT DNA detection by qPCR

For the purification of genomic DNA from WJ-MSC273 and WJ-MSC/TERT273 cells and from the respective secreted EVs, the QIAamp® DNA Micro Kit (50) (Qiagen) was used. Minor adaptations to the protocol for genomic DNA isolation from small volumes of blood were made due to the different nature of the samples. WJ-MSC273 and WJ-MSC/TERT273 cells were collected, and the cell pellets (3 x 10^5^ cells) were washed twice with 500 μl of PBS after centrifugation for 5 minutes at 300 x g. The cell pellets were then resuspended in 50 μl of PBS before adding 50 µL of buffer ATL to reach a final volume of 100 µL / sample. Then, 10 µL proteinase K and 100 µL of Buffer AL were added to each sample before mixing by pulse-vortexing for 15 seconds. The homogeneous solution was incubated at 56°C for 10 minutes at 550 rpm to increase DNA extraction yield. All the following steps were performed according to the manufacturer’s protocol, and the final elution step was performed in 20µL of HyPure^TM^ molecular biology grade water (Cytiva). Similarly, the purification of genomic DNA from WJ-MSC273 and WJ-MSC/TERT273-derived EVs was performed by adding 50 µL of buffer ATL to 50µL of EV-containing samples (2 - 5.5 x 10^9^ particles per sample) to reach a final volume of 100 µL / sample. The EV-containing samples were then processed as described above and the final elution step was performed in 20µL of HyPure^TM^ molecular biology grade water (Cytiva). DNA quantification was performed by Thermo NanoDrop One ND-ONE W Mikrovolumen-UV/VIS S (Thermo Fisher Scientific). A minimum value of A260/A280 ratio of 1.70 was used as threshold for DNA purity of the DNA samples. The obtained DNA was used for the set-up of qPCRs for the detection of hTERT DNA in WJ-MSC273 and WJ-MSC/TERT273 cells and from the secreted EVs. A master mix was obtained by mixing 2.2 µL of HyPure^TM^ molecular biology grade water (Cytiva), 0,8 µL primer mix (5 µM Forward primer and 5 µmM Reverse primer) and 5 µL of 2X TATAA SYBR GrandMaster Mix 1875 rxn (TATAA), for a final volume of 8 µL / well of master mix to be pipetted into the bottom of each well of 96-Well-PCR-Plates (Starlab). Then, 2 µL of pre-diluted DNA was pipetted directly into the master mix and mixed by gentle pipetting. Cell-derived DNA and EV-derived DNA were diluted according to the lower DNA concentration obtained from each DNA extraction from EV-containing samples; cell-derived DNA was also tested at fixed concentrations of 3.7 ng/µL and 10 ng/µL. For each experiment, hTERT-carrying plasmid DNA was used as positive control, while molecular biology grade water (Cytiva) was used as negative control. Xtra-Clear Advanced Polyolefin StarSeal (qPCR) (Starlab) was used to seal the plate before spinning at 700 x g for 30 seconds. The LightCycler®96 (LC96) instrument (Roche) was used to read the plate, and a standard 3-step amplification program (denaturation for 5 seconds at 95°C, annealing at 58°C for 30 seconds and extension at 72°C for 10 seconds) was run for 35 cycles. LC96 software was used for data analysis. Both the Forward (sequence: GAGAACAAGCTGTTTGCGGG) and Reverse (sequence: AAGTTCACCACGCAGCCATA) primers for hTERT detection (Invitrogen) were designed to align to exon 9 of genomic hTERT, with resulting amplicon size of 151 bp.

### 2.4 Analysis of the biological activity of EV preparations

#### 2.4.1 Anti-inflammatory Assay

To test the anti-inflammatory properties of WJ-MSC273 and WJ-MSC/TERT273-derived EVs, 4 x 10^4^ RAW 264.7 cells/well were seeded in serum-containig cell culture medium (see 2.1.1) in a 96-well plate. Cells were incubated at 37°C (5% CO_2_, ambient oxygen) for 24 hours prior to treatment. All treatments were prepared in starvation medium (Minimum Essential Medium Eagle, Sigma) supplemented with GlutaMAX^TM^-I (Gibco), with a final volume of 100 µL/well. For each treatment, RAW 264.7 cells were tested in parallel with and without stimulation with 100 ng/mL of lipopolysaccharide (LPS) from E. coli 0127:B8 (Sigma) in triplicates. Before treatment, cells were washed with 80 µL of starvation medium. Thereafter, TFF-enriched 1 x 10^9^ particles/mL from WJ-MSC273 and WJ-MSC/TERT273 cells were added to each well in triplicates. 20 mM HEPES and TFF enriched cell-free EV collection medium were included as negative controls. As positive control, 2.5 µMmL Dexamethasone (Sigma, 1mM dissolved in DMEM-Ham’s F12 1:1) was included. Total volume of 20 mM HEPES was kept constant across all treatment conditions. 24 hours after the treatment, the NO secretion was measured using Griess Reagent (Promega) following the manufacturer’s instructions. After 5 minutes of light-protected incubation with Sulfanilamide at RT and 5 minutes with N-1-napthylethylenediamine dihydrochloride (NED Solution), absorbance was measured at 535 nm with a Spark^®^ multimode microplate reader (Tecan).

#### 2.4.2 Anti-fibrosis Assay

To assess the anti-fibrotic properties of WJ-MSC273 and WJ-MSC/TERT273-derived EVs, 2.5 x 10^3^ fHDF/TERT166 cells/well were seeded into a 96 well plate in starvation medium (DMEM/Ham’s F-12 1:1 supplemented with 0.5% FBS, Primocin (InvivoGen) and 100 μg/mL G418). Cells were incubated at 37°C for 24 hours prior to treatment. All treatments were prepared in starvation medium and performed in sextuplicates. In all treatments - with the only exception of negative controls – alpha smooth muscle actin (α-SMA) induction was induced by adding 100 pg/mL of rhTGFbeta-1 (Abcam) to each well. Treatments included 1 x 10^8^ or 1 x 10^9^ EVs/mL. As negative control, 20 mM HEPES or TFF enriched medium were used. 2 μM PP2 (Avantor/VWR, stock diluted 1:25) was used as a positive control. The total volume of 20 mM HEPES was kept constant across all treatment conditions. The treatment was repeated on day 2 and 5 after the first treatment. The plate was incubated at 37°C until α-SMA induction analysis. 24 hours after the last treatment, cells were washed twice with PBS Ca^2+^/Mg^2+^, fixed with Roti-Histofix (4% paraformaldehyde, Carl Roth) for 10 minutes at RT and washed again three times with PBS Ca^2+^/Mg^2^. Then, cells were permeabilized by adding blocking solution (5% BSA/0.2% Triton-X-100 (Sigma) in PBS Ca^2+^/Mg^2^) and incubating for 20 minutes at RT. The following primary antibodies were used for cell staining overnight at 4°C: anti-alpha smooth muscle actin [1A4] antibody (Abcam) and isotype control mouse IgG2a (Sigma) diluted 1:200 and 1:40 respectively in blocking solution (5% BSA in PBS, Sigma).. After two washes with PBS Ca^2+^/Mg^2^, cells were stained with donkey anti-mouse AF594 secondary antibody (Jackson, 1:500) in PBS Ca^2+/^Mg^2+^. Finally, the plate was washed twice with PBS Ca^2+^/Mg^2^ and α-SMA induction was measured on a Spark® multimode microplate reader (Tecan) with an excitation at 580 nm and an emission at 615 nm.

#### 2.4.3 Wound Healing Assay

To investigate the wound healing properties of WJ-MSC273 and WJ-MSC/TERT273-derived EVs, Culture-Insert 2 Well in µ-Dish 35 mm (ibidi) were carefully placed into wells of a 24-well plate. HUVEC/TERT2 cells were harvested and 75.000 cells/cm^2^ (equivalent to 16.500 cells/insert, resuspended in complete culture medium) were seeded into the inserts. Cells were incubated at 37°C for 24 hours prior to treatment. Inserts were carefully removed, and the wells were washed twice with PBS before addition of 250 µL of starvation medium/well (EBM™ basal medium supplemented with hydrocortisone, hEGF, ascorbic acid, BBE and G418 at the same concentration as in complete medium) for the imaging of the gaps at time 0. For all treatments, with the exception of the positive control, for which complete culture medium was used, starvation medium was added to each well to a final volume of 500 µL/well. Cells were treated with 1 x 10^8^ or 1 x 10^9^ EVs/mL. As negative controls, TFF-enriched cell-free EV production medium and 20 mM HEPES were used. The volume of 20 mM HEPES was kept constant across all treatments to 1 mM per well. The imaging of the gaps was done after the removal of the inserts (time 0) and 16 and 40 hours after treatment using an EVOS M5000 microscope. For each well, 5 pictures of the gap between the inserts were taken, while the free-hand tool of EVOSM5000 was used to draw the edges of the gaps and to calculate the free area between the wells of the inserts. The percentage of gap closure was calculated by normalizing the free area after 16 and 40 hours post-treatment with the free area at time 0.

### 2.5 Statistical analysis

GraphPad Prism 9.0 (GraphPad Software, Inc. San Diego, CA, USA) was used for all the statistical analysis. At least three biological replicates have been analyzed for each assay with triplicate technical replicates. Bars represent mean ± SD. Direct comparison of WJ-MSC273 and WJ-MSC/TERT273 differentiation capacity and the characterization of the respective secreted EVs was analyzed with an unpaired t-test. The biological activity of primary and TERT EVs was analyzed with two-way ANOVA unless mentioned differently. Significance levels are reported as follows: * 0.01 ≤ p < 0.05; ** 0.001 ≤ p < 0.01; *** 0.001 ≤ p < 0.0001; and **** p < 0.0001.

## RESULTS

### 3.1 hTERT mediated immortalization does not affect Wharton’s Jelly mesenchymal stromal cell morphology and growth properties

To investigate whether ectopic expression of hTERT alters the cellular phenotype, WJ-MSC/TERT273 telomerized cells were compared to the non-transfected parental primary cells. First, growth characteristics and morphology were investigated. Late-passage WJ-MSC/TERT273 cells maintained a morphology that was very similar to early-passage cells, while WJ-MSC273 primary cells at late population doublings (PDs) showed morphological changes typical of replicative senescence, with enlarged, flattened cells (Fig. 1A). In terms of cell proliferation, primary and hTERT immortalized WJ-MSCs displayed similar growth rates of 1.7–1.8 PDs/day corresponding to a population doubling time of about 14 hours (Fig. 1B). Notably, while primary cell growth rates declined at the end of the replicative life span, WJ-MSC/TERT273 cells were continuously cultivated for more than 110 PDs with no signs of changed proliferation rates (Fig. 1C). This life span extension is almost twice as long as the one of the primary cells, which showed marked slowing of the growth rate after 65.5 PDs growth arrest after 71.9 PDs. In line with decreased cell growth, late-passage WJ-MSC273 cells showed increased SA-β-Galactosidase activity as a marker for replicative senescence when compared to early-passage primary cells (Sup. Fig. 1A, 1B), while hTERT immortalized WJ-MSCs at early- and late-passage showed similarly low SA-β-Galactosidase activity as early-passage primary WJ-MSCs. To confirm that hTERT transfection indeed resulted in induction of telomerase activity (TA), the RT-TRAP and TeloTAGGG Telomerase PCR ELISA^plus^ assays were used to measure relative TA normalized to a commercial standard or to HEK293 cells. While WJ-MSC273 showed TA just above the detection limit, WJ-MSC/TERT273 cells displayed clear telomerase activity (Fig. 1D). This was corroborated at early and late PDs (Sup. Fig. 1C).

**Figure 1.**
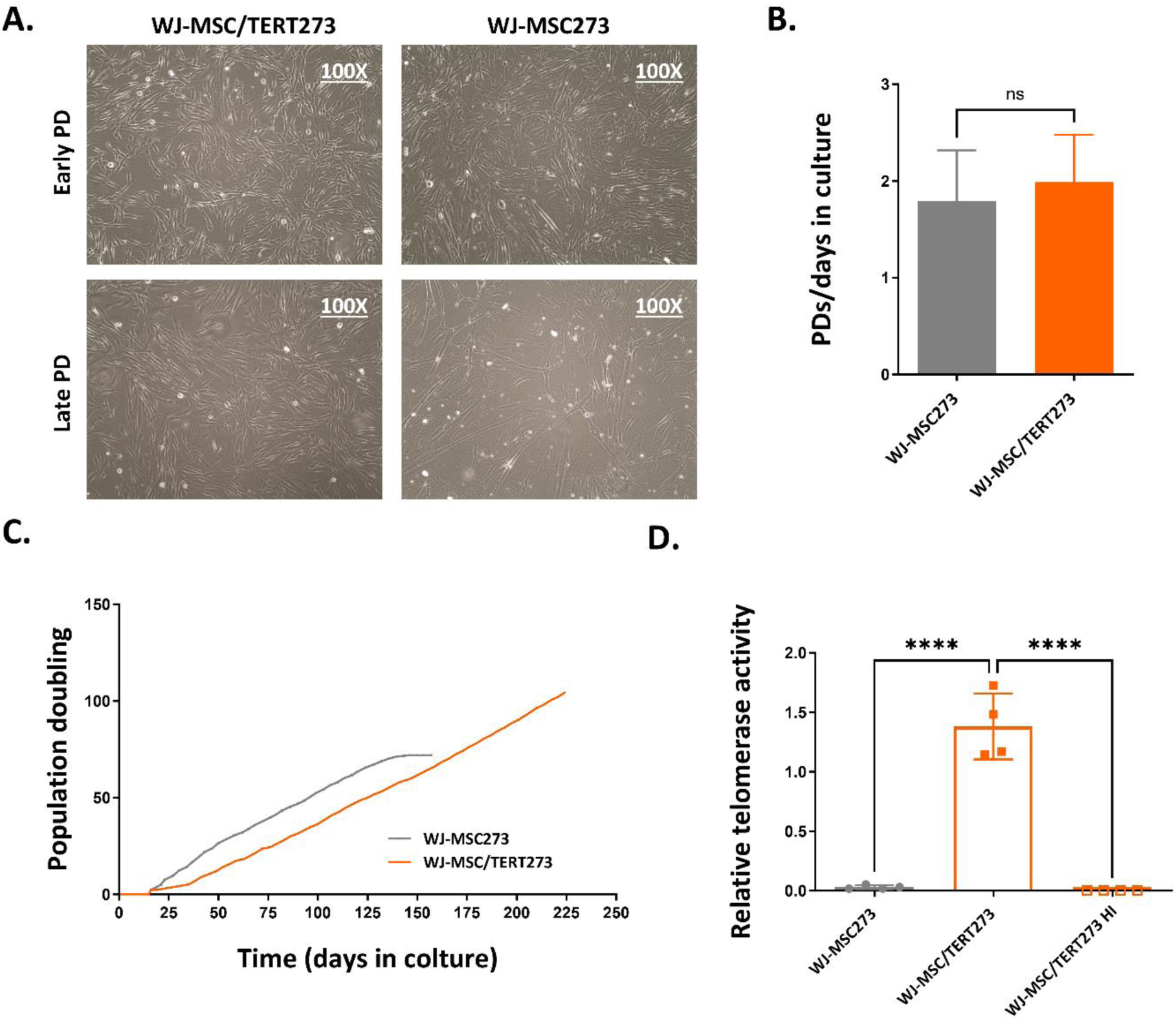
Comparison of cell morphology, growth potential and telomerase activity between primary and telomerized WJ-MSCs. **A.** Morphological comparison of primary and telomerized WJ-MSCs at early (PD14.2 and 14.8) and late population doubling times (PD71.4 and 76.0) by bright field microscopy. Images were taken at 100X magnification. **B.** Mean PD rate of primary and telomerized WJ-MSCs during exponential growth (from day 16 to day 134, n=35). **C.** Growth curves of primary and telomerized WJ-MSC during long-term culture. The cumulative population doublings (PDs) are reported as a function of the culturing days and measured at each passage. **D.** Evaluation of telomerase activity in primary and telomerized WJ-MSCs by Telomere Repeat Amplification Protocol (TRAP) assay at PD18.6 and PD24.4 respectively (n=4). Telomerase activity in WJ-MSC/TERT273 upon Heat Inactivation (HI) was measured as negative control. Significance levels are reported as follows: **** p < 0.0001. The data are presented as mean ± SD.

### 3.2 hTERT mediated immortalization does not affect Wharton’s Jelly mesenchymal stromal cells functional properties

We next assessed the expression of typical MSC surface markers. Flow cytometric analysis confirmed the presence of MSC-specific markers CD105, CD90, CD73, and the absence of CD34 at similar levels in WJ-MSC273 cells and their immortalized counterpart (Fig. 2A). To evaluate MSC functionality following hTERT overexpression, we compared the tri-lineage differentiation capacity of primary and immortalized WJ-MSCs. First, osteogenic differentiation of primary and TERT WJ-MSCs was induced and analyzed after 21 days. Both WJ-MSC273 and WJ-MSC/TERT273 displayed the expected alterations in cellular morphology and stained positive for extracellular calcium deposits using Alizarin red (Fig. 2B), with no significant differences (Fig. 2C). Next, we examined the adipogenic differentiation capability of WJ-MSC273 and its corresponding immortalized counterpart. Oil Red-O staining of lipid droplets was used to confirm adipogenic differentiation three weeks after induction (Fig. 2D) and was quantified by absorbance measurement at 520 nm, again showing no significant differences (Fig. 2E). Finally, WJ-MSCs were tested for chondrogenic differentiation. Following two days of growth in conventional tubes in chondrogenic differentiation medium the cell pellets lost adhesion and formed spherical aggregates that grew over time, indicating the development of extracellular matrix. After 21 days of differentiation, cell pellets were stained with Alcian blue to mark the glycosaminoglycans in cartilage. Both primary and immortalized WJ-MSCs exhibited chondrogenic differentiation characteristics, as evidenced by similar dark-blue staining of the extracellular cartilage matrix and by the presence of isogenous chondrocyte groups scattered throughout the matrix (Fig. 2F). Interestingly, WJ-MSC273 pellets displayed a more stratified cartilaginous matrix at the edges and a denser chondron growth at the core, while the pellets from WJ-MSC/TERT273 cells showed the opposite orientation (Fig. 2F). This variability in cartilaginous matrix formation is often observed when differentiation is performed under normoxic conditions (Lee et al., 2013; Leijten et al., 2014). These results suggest that both cell strains can undergo chondrogenesis, but there are differences in its efficiency. Summarized, these data indicate that WJ-MSC/TERT273 cells maintain both MSC-typical surface marker expression and trilineage differentiation potential comparable to their primary counterpart during expansion under standard culture conditions.

**Figure 2.**
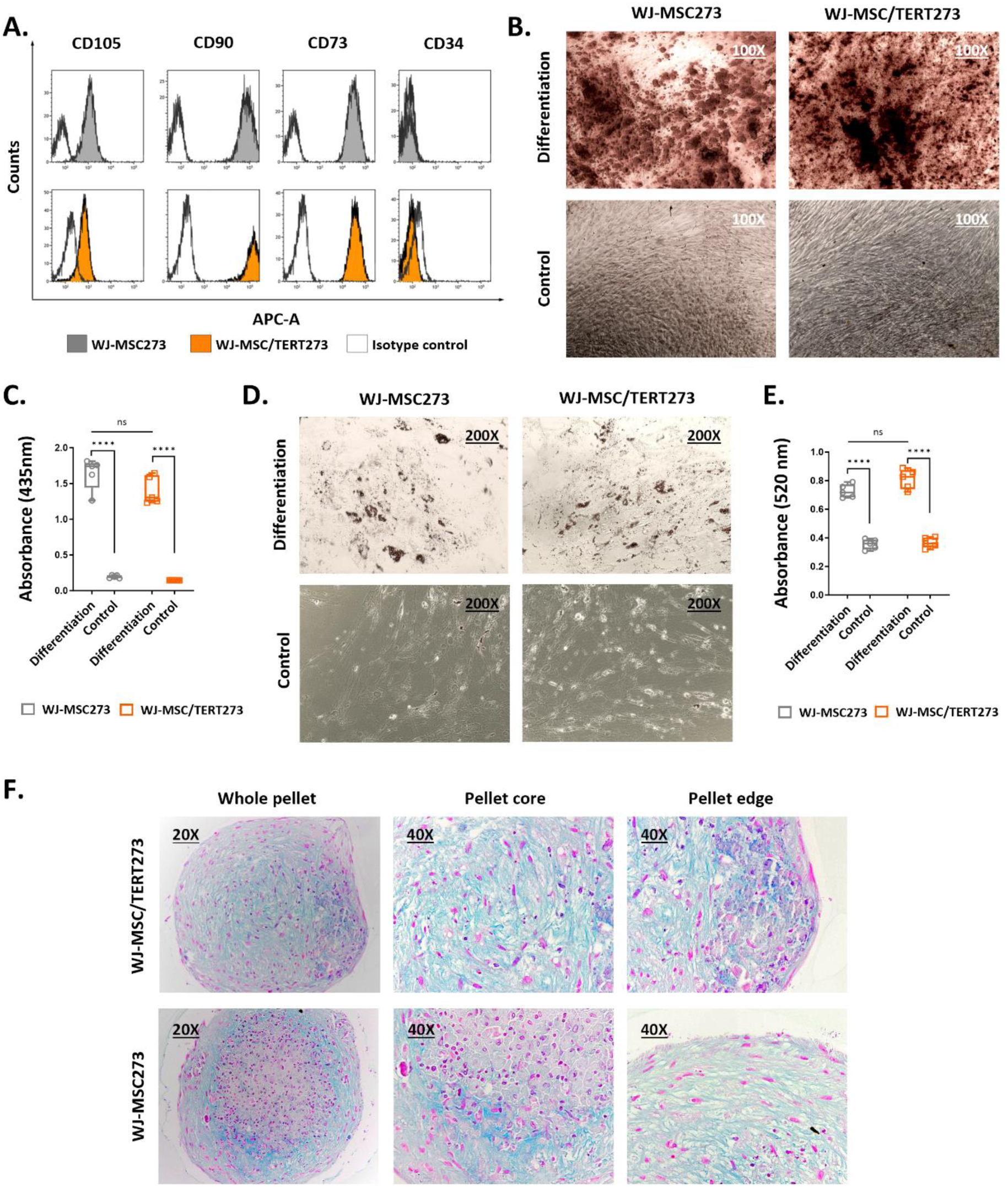
Comparison of surface marker expression and tri-lineage differentiation potential between primary and telomerized WJ-MSCs. **A.** Evaluation of canonical MSC surface marker expression in primary and telomerized WJ-MSCs by flow cytometric analysis. Marker expression by primary WJ-MSCs is shown in gray, telomerized WJ-MSCs in orange, and isotype control in white. **B.** Representative microscopy images of Alizarin red staining show calcium deposition in primary and telomerized WJ-MSCs upon osteogenic differentiation induction (100X magnification) **C.** Quantitation of osteogenic differentiation potential by quantification of Alizarin red staining in primary and telomerized WJ-MSCs. **D.** Representative microscopy images of Oil Red O staining show lipid droplets in primary and telomerized WJ-MSCs after adipogenic differentiation (100X magnification). **E.** Quantitation of adipogenic differentiation potential by quantification of Oil Red O staining in primary and telomerized WJ-MSCs. **F.** Representative microscopy images of Alcian blue staining show acidic mucopolysaccharide and sialomucin substances in primary and telomerized WJ-MSCs upon chondrogenic differentiation induction (left panel 20X; middle and right panels 40X magnification). Quantitative data are presented as mean ± SD of the absolute absorbance values at respective wave lengths; n=5 for differentiation conditions and n=3 for controls. Significance levels are reported as follows: **** p < 0.0001.

### 3.3 Characterization of primary and telomerized WJ-MSC-derived EVs

To investigate whether hTERT-mediated immortalization of WJ-MSCs influences the secreted EVs, we isolated WJ-MSC273 and WJ-MSC/TERT273-derived EVs from conditioned medium (CM) of cells that had been cultured in serum-free medium for 48 hours. After two low-speed centrifugations and a 0.22 µm filtering step, EVs were enriched from the collected CM by TFF using a 300kDa cut-off column before characterization (Fig. 3A). NTA analysis demonstrated that the concentration and particle size distribution of isolated EVs from 3 consecutive batches were comparable in size and number between primary and immortalized cells (representative image Fig. 3B). Both primary and immortalized WJ-MSCs secreted 1.1×10^4^ - 1.7×10^4^ particles per cell per day (Fig. 3C), which is in the range reported in various other studies (Ragni et al., 2020; Walravens et al., 2021) As controls, the TFF-enriched cell-free medium was also analyzed by NTA and compared to both EV preparations, it showed a 7-to-11-fold lower particle concentration (Sup. Fig. 2A). Protein concentrations in EV preparations were assessed using the BCA assay, and the resulting data were normalized to particle counts (Fig. 3D) showing similar amounts of protein / particle. The particle mean diameter (Fig. 3E) was in line with the expected EV size range (ranging from 140 to 172 nm) and consistent between primary WJ-MSC EVs and their telomerized counterparts. To note, the size of particles in the EV collection medium was significantly lower compared to both primary and telomerized WJ-MSC EVs (Sup. Fig. 2B) as visualized by NTA measurement (Sup. Fig. 2C). Moreover, we used transmission electronic microscopy (TEM) to assess the morphology and size of primary and TERT EVs, ranging between 100-200 nm (Fig. 3F).

**Figure 3.**
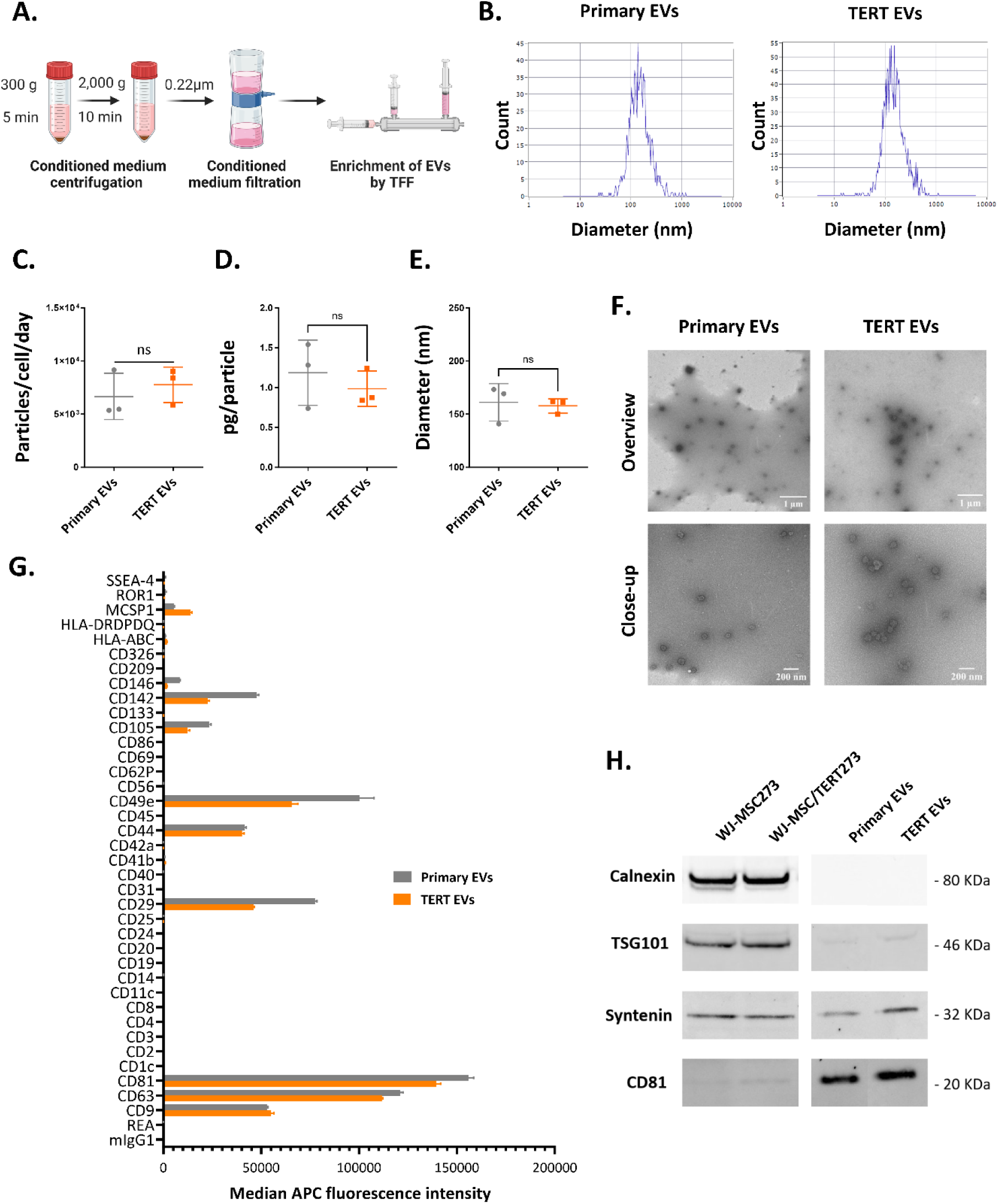
Characterization of primary and telomerized WJ-MSC derived EVs. **A.** Schematic representation of the experimental set-up used for the enrichment of primary and telomerized WJ-MSCs-derived EVs. The conditioned media from primary and telomerized WJ-MSCs are collected, centrifugated and filtrated prior to EVs enrichment though 300kDa Tangential Flow Filtration (TFF). **B.** Representative size distributions of primary (left) and telomerized (right) WJ-MSC derived EVs by Nanoparticle Tracking Analysis (NTA). **C.** Particle release from primary and telomerized WJ-MSCs by NTA. Results are normalized to the cell numbers and time of EV secretion. **D.** Protein concentration of the TFF-enriched EVs from primary and telomerized WJ-MSCs by BCA analysis. **E.** Mean particle diameter from primary and telomerized WJ-MSCs by NTA. For particle release, particle size and protein concentration of EVs, data are presented as mean ± SD (n=3). **F.** Transmission electron micrography (TEM) of EVs isolated from primary and telomerized WJ-MSCs. Scale bar =500 nm. **G.** MACSPlex assay of the TFF-enriched EVs from primary and telomerized WJ-MSCs. The mean of median APC fluorescence intensities is shown on y-axis (n = 3) for all markers included in the kit (x-axis). **H.** Western Blot analysis of CD81, Syntenin-1, TSG101 and Calnexin for cell lysate from primary and telomerized WJ-MSCs and of the respective tetraspanin-positive primary and TERT EVs.

EV surface marker profiles were assessed by multiplex bead-based flow cytometry. Very similar surface protein profiles for CD9, CD63, CD81, and CD44 on primary and TERT EVs were observed with slightly lower presence of CD142, CD105, CD29 and CD49e on TERT EVs (Fig. 3G). We further confirmed the enrichment of the EV preparations by assessing the absence of Calnexin and the presence of TSG101, CD81 and Syntenin by Western Blot analysis (Fig. 3H) in tetraspanin positive EVs. Using human CD81+ bead-based flow cytometry analysis, we confirmed the presence of CD9, CD81, and CD63 on both EV types (Sup Fig. 3A, B).

### 3.4 Small RNA analysis reveals comparable miRNA expression profiles in primary and telomerized WJ-MSC-derived EVs

To analyze the impact of hTERT immortalization on the miRNA cargo of the released particles, we isolated total RNA from 1 x 10^9^ primary and TERT EVs. SmallRNA sequencing was performed according to the experimental workflow described in Sup. Fig. 4A resulting in between 60 and 80 million raw reads per sample. The absolute read counts of WJ-MSC273 and WJ-MSC/TERT273 EV samples (2.8 x 10^7^ and 2.6 x 10^7^ total reads, respectively) consisting of various RNA-types, unclassified genomic, and unmapped reads, were comparable (Fig. 4A). Additionally, WJ-MSC273- and WJ-MSC/TERT273-EVs had similar absolute miRNA read counts for processed reads examined using miRDeep2 between EVs from WJ-MSC273 and WJ-MSC/TERT273 (Fig. 4B). Considering that the more abundant miRNAs are most probably contributing most to the biological activity of EVs, the overlap between the top 20% abundant miRNAs based on the average rpm expression was investigated. Of the 76 miRNAs in WJ- MSC273-EVs and 69 miRNAs in WJ-MSC/TERT273-EVs representing the top 20% abundant ones, 65 miRNAs were found to be shared among the two groups (Fig. 4C). This accordance of expressed miRNAs was also observed in a correlation analysis over all detected miRNAs with a correlation coefficient of 0.82 and a p-value < 0.001 (Fig. 4D). Still, some differentially expressed miRNAs were identified when comparing the miRNome profiles of WJ-MSC273- and WJ-MSC/TERT273-EVs with 8 out of the 214 FDR based filtered miRNAs showing significantly altered expression with an FDR < 0.05. In primary WJ-MSC EVs, 4 miRNAs were found to be more and 4 less abundant, whereas 2 of the high abundant miRNAs were found with a logFC > 2 (Sup. Fig. 4B, C). Overall, these data suggest that primary and telomerized WJ-MSC-derived EVs have comparable, albeit slightly different miRNA expression profiles.

**Figure 4.**
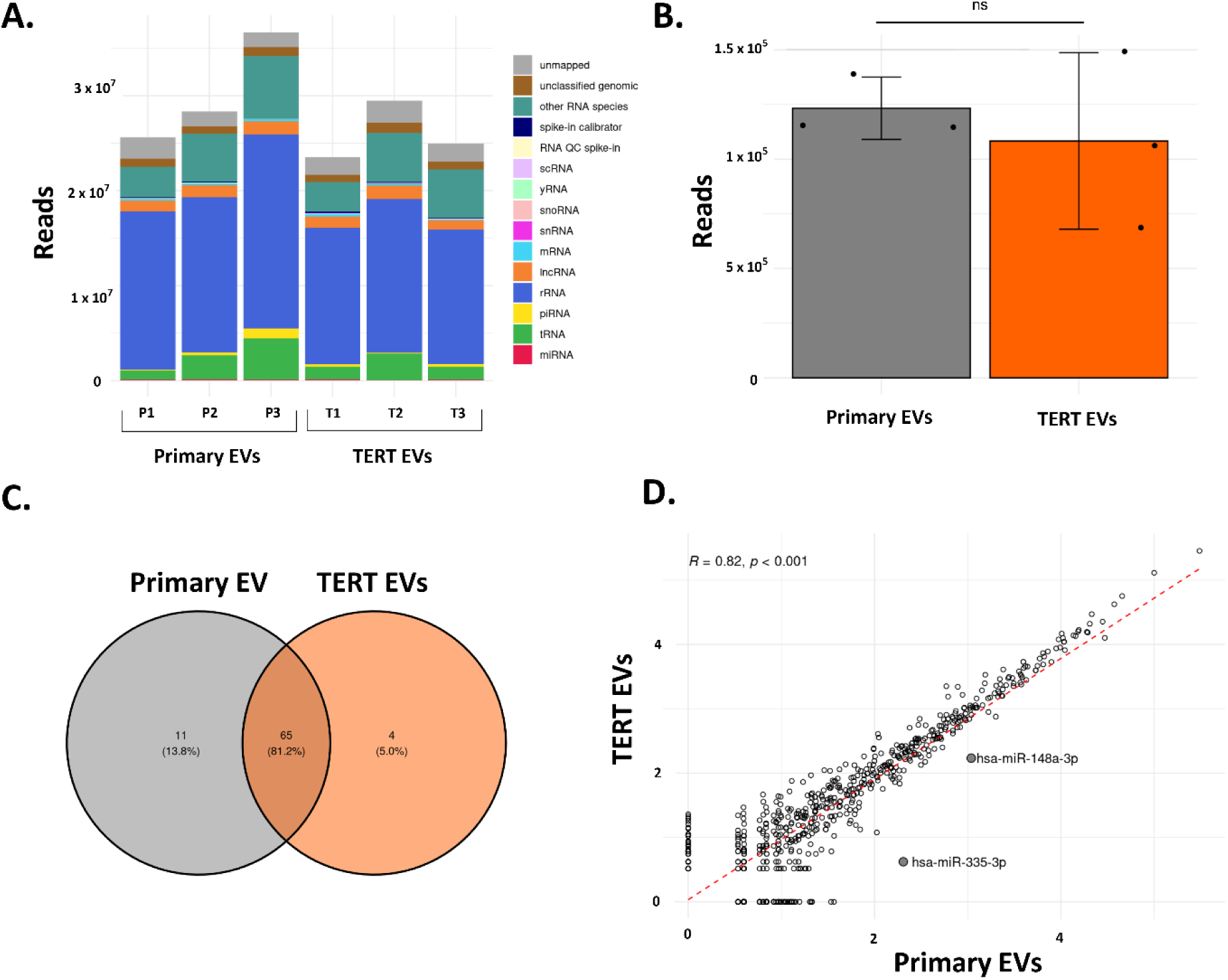
Differential expression of microRNAs in primary and telomerized WJ-MSC derived EVs. **A.** Absolute read counts of different RNA-types, unclassified genomic and unmapped reads. **B.** Normalized number of mapped reads (RPM) used for bowtie mapping of small RNA sequencing reads against the gene sequence of human telomerase. **C.** Venn diagram of the overlap of the top 20% expressed miRNAs in primary and TERT EVs. Expression (rpm) was filtered for the detection of miRNAs in all 3 replicates of either TERT or primary EVs. Mean rpm was calculated and ranked accordingly. Only the top miRNAs representing 20% of the highest rpm abundance were selected and compared in the Venn diagram. **D.** Differentially expressed miRNAs between primary and hTERT-MSC EV-cargo. Expression correlation of the gene expression is visualized using a robust linear model, and miRNAs with a False Discovery Rate (FDR) <0.01 and absolute logarithmic Fold Change value (logFC) > 2 are displayed and highlighted. Spearman rank test is used to calculate the correlation coefficient and significance

### 3.5 Primary and telomerized WJ-MSC-derived EVs show similarities in their biological activity in vitro

To exclude that hTERT-mediated immortalization of WJ-MSCs affects the bioactivity of the secreted EVs, we compared the biological activities of primary and TERT EVs using different cell-based assays. First, we investigated the anti-inflammatory potential of WJ-MSC273 and WJ-MSC/TERT273 EVs. LPS-stimulated RAW 264.7 cells were treated with 1 x 10^9^ particles/mL and 24 hours after treatment, Nitric Oxide (NO) production was measured in the culture medium. EVs from both cell sources showed 40 to 50% reduction in NO secretion compared to both vehicle and medium controls, using three independent EV enrichments (Fig. 5A). Then, we compared the anti-fibrotic activity of WJ-MSC273 and WJ-MSC/TERT273 EVs. Therefore, telomerized human dermal skin fibroblasts (fHDF/TERT166) were exposed to TGF-β as an inducer of fibrosis and treated with EVs or the kinase inhibitor PP2 as a positive control. As read-out, alpha smooth muscle actin (α-SMA) staining using immunofluorescence microscopy was used. Both primary and telomerized EVs reduced α-SMA expression by 25% in TGFβ1-stimulated fHDF/TERT166 compared to both vehicle and medium control at 1 x 10^9^ particles/mL (Fig. 5B). Since α-SMA expression is a driving event in pro-fibrotic tissue repair, these data suggest that both EVs from primary and telomerized MSCs might exert similar anti-fibrotic activityFinally, we investigated the effect of EVs from both primary and telomerized WJ-MSCs on cell motility and migration of telomerized human umbilical vein endothelial cells (HUVEC/TERT2) using a scratch assay as a surrogate test for wound healing. After removal of an insert generating reproducible ‘wounds’ in the cell layers, we incubated the cells with serum-free medium supplemented with WJ-MSC/TERT273 and WJ-MSC273 EVs. 10% serum-containing medium was used as a positive control. 40-hours post-treatment a dose-dependent decrease of the open scratch was observed (Fig. 5C, D, Sup. Fig. 5A), with no significant differences between cell sources. Taken together, these results suggest a highly similar bioactivity profile of EVs from primary and telomerized MSCs.

**Figure 5.**
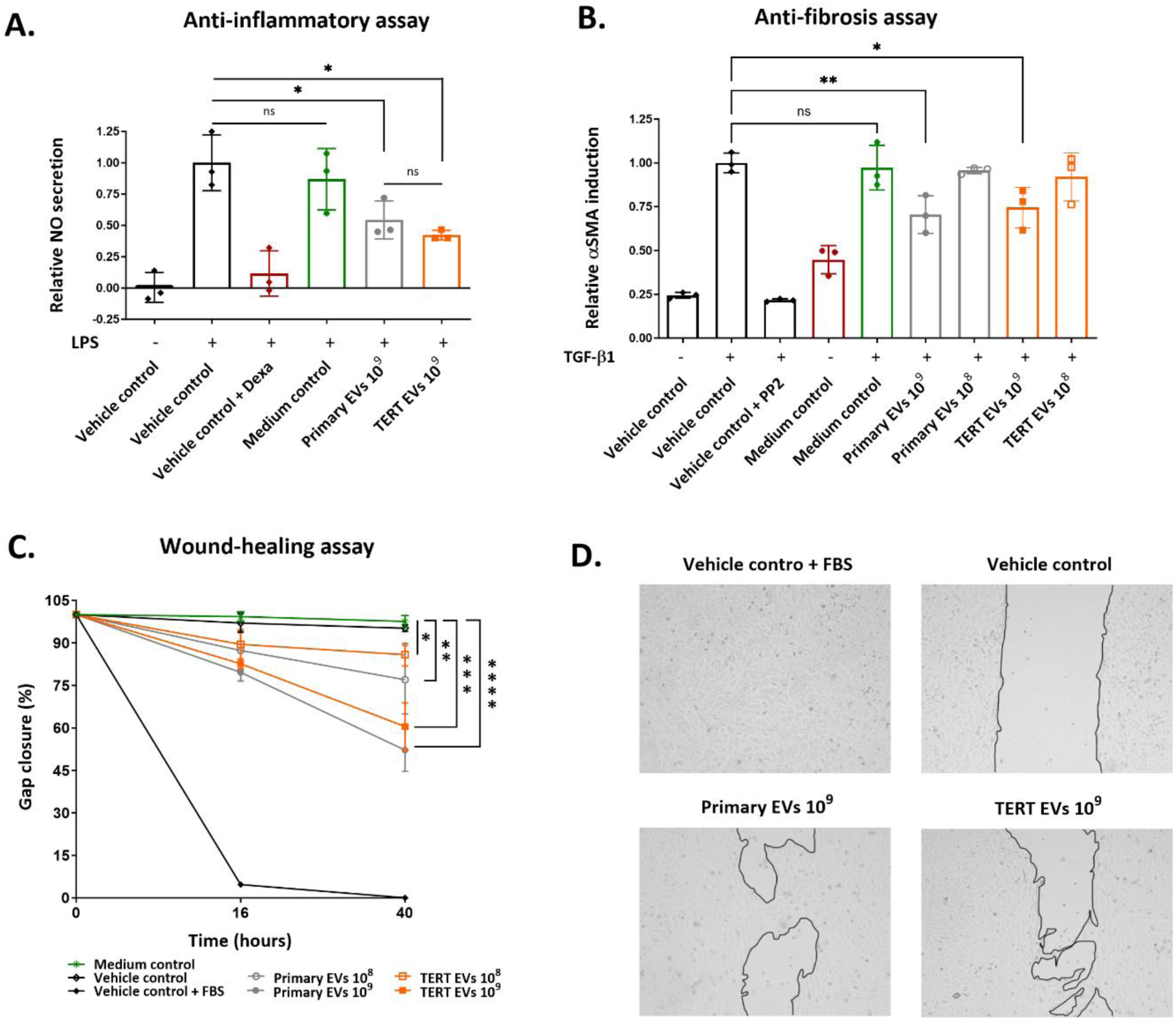
Evaluation of the biological properties of primary and telomerized WJ-MSC derived EVs. **A.** Evaluation of the anti-inflammatory activity of primary and TERT EVs on nitric oxide (NO) secretion by LPS-stimulated RAW264.7 cells. 20 mM HEPES (vehicle control) and TFF enriched cell-free EV collection medium (medium control) were included as negative controls, while 2.5 µMmL Dexamethasone (vehicle control + Dexa) was included as positive control. **B.** Evaluation of the anti-fibrotic activity of primary and TERT EVs on α-SMA induction by TGF-β1-stimulated fHDF/TERT166 cells. 20 mM HEPES (vehicle control) and TFF enriched cell-free EV collection medium (medium control) were included as negative controls, while 2 μM PP2 (vehicle control + PP2) was included as positive control. **C.** Quantification of the wound closure of HUVEC/TERT2 cells at 0, 16 and 40 h after the removal of culture insert and treatment. 20 mM HEPES (vehicle control) and TFF enriched cell-free EV collection medium (medium control) were included as negative controls, while complete culture medium (vehicle control + FBS) was included as positive control. Significance levels are reported as follows: * 0.01 ≤ p < 0.05; ** 0.001 ≤ p < 0.01; *** 0.001 ≤ p < 0.0001; **** p < 0.0001. The data are presented as mean ± SD, with n=3 for each assay. **D.** Representative microscopy images of the wound closure of HUVEC/TERT2 cells 40 h after the removal of culture insert and treatment.

### 3.6 Overexpression of hTERT neither increases hTERT in EVs nor induces anchorage independent proliferation of MSCs

We next sought to investigate whether hTERT-mediated immortalization of WJ-MSC273 could result in the presence of hTERT mRNA in telomerized MSC-EVs. Therefore, we assessed the number of NGS reads generated above (Fig. 4A) that would map to the human telomerase genomic regions (Fig. 6A) and mRNA transcript sequences (Sup. Fig 6A). The average size distribution of mapped reads showed a comparable distribution for primary and telomerized MSC-EVs (Sup. Fig 6A). Additionally, the normalized number of aligned reads did not show a significant difference between the two groups (Fig. 6A). For the specific mapping against the transcript sequence (NM_198253.3), only a few hits for telomerized MSC-EV samples with RPM values not higher than 0.6 were found (Sup. Fig. 6B). Despite this low transcript level - usually reads with such low RPM values are filtered out - we investigated location and length of the reads using Integrated Genomic Viewer. Coverage plots (Sup. Fig. 6C) showed no specific mapping regions against the gene sequence of hTERT, with reads mapping to the gene sequence mainly associated with intronic regions and comparable between primary and TERT EVs. Such a low mapping rate, together with the investigation of coverage plots showing no specific mapping regions against the coding sequence of hTERT suggest mainly random hits for the TERT transcript that do not differ between telomerized and primary MSCs.

**Figure 6.**
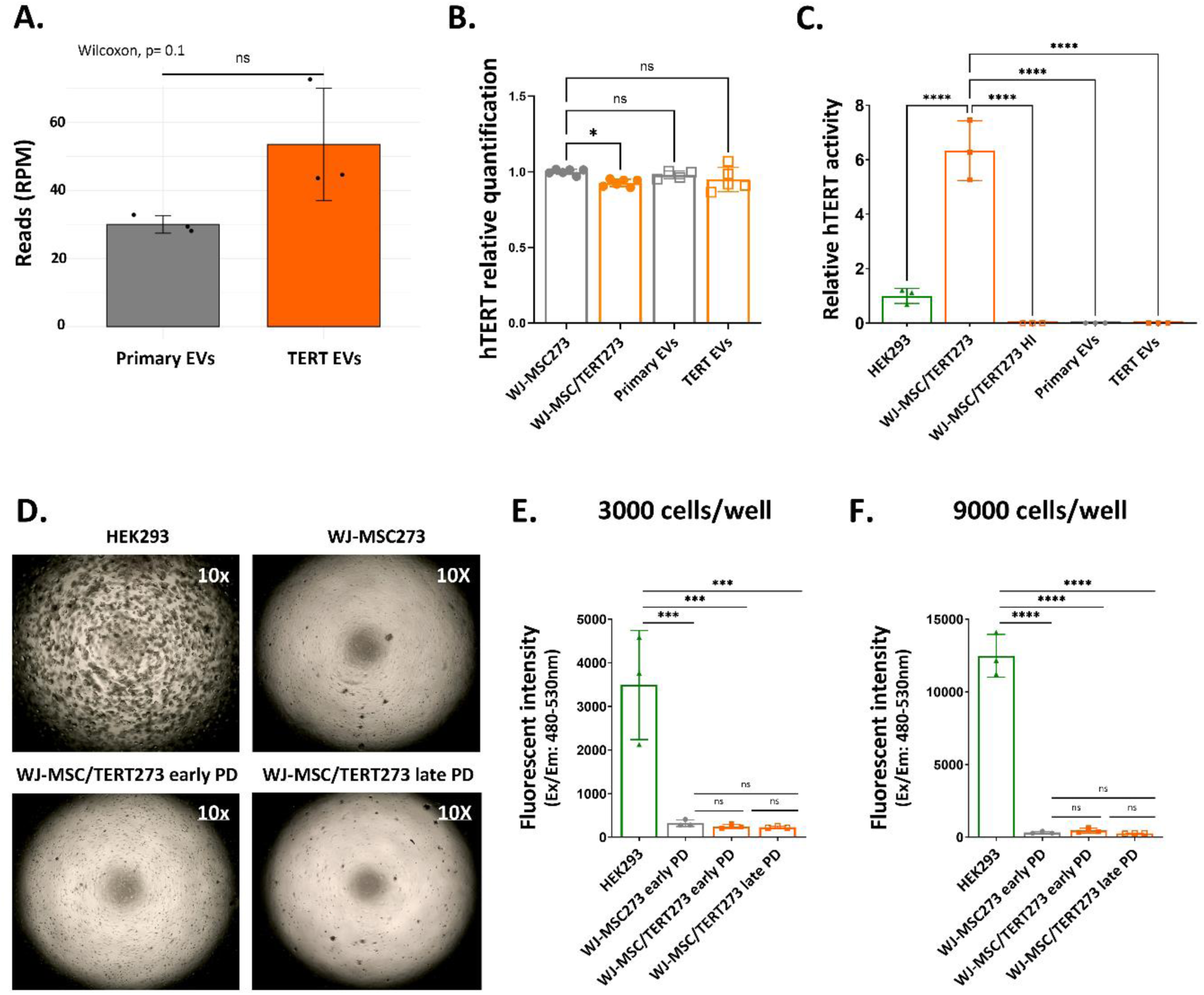
Evaluation of the safety profiles of primary and telomerized WJ-MSCs and the respective EVs. **A.** Normalized number of mapped small RNA sequencing reads against human telomerase. **B.** Quantification by qPCR of hTERT genomic DNA in WJ-MSC273 and WJ-MSC/TERT273 cells, and in the respective secreted EVs. Results are normalized to the ct values of WJ-MSC273 primary cells. **C.** Relative telomerase activity in telomerized WJ-MSCs and EVs enriched from primary and telomerized WJ-MSCs using the TeloTAGGG Telomerase PCR ELISAPLUS assay. WJ-MSC/TERT273 cells were tested at PD 58.4; the hTERT activity was calculated relative to HEK273 cells, which were measured as positive control. Heat Inactivation (HI) was performed on WJ-MSC/TER273 cells as negative control. **D.** Assessment of primary and telomerized WJ-MSCs tumorigenicity at different population doubling time by colony formation in soft-agar. Representative microscopic images are shown, and HEK293 cells were used as positive control for tumorigenesis. Images were taken at 10X magnification **E-F.** Quantification of primary and telomerized WJ-MSCs tumorigenicity by fluorescent intensity measurement. Cells were seeded at a density of 3000 and 9000 cells per well respectively. Significance levels are reported as follows: * 0.01 ≤ p < 0.05; *** 0.001 < 0.0001; and **** p < 0.0001. The data are presented as mean ± SD (n=6) for qPCR data of primary and telomerized WJ-MSCs, as mean ± SD (n=5) for qPCR data of primary and TERT EVs and as mean ± SD (n=3) for TeloTAGGG Telomerase PCR ELISAPLUS assay and for colony formation assay in soft-agar.

Moreover, we verified the lack of enrichment of hTERT DNA in primary and TERT EVs by qPCR (Fig. 6B and Sup. Fig. 7A, B). In more detail, a qPCR without reverse transcription step was performed using forward and reverse primers matching to exon 9 of hTERT DNA to assess hTERT DNA amplification. DNA extraction resulted in similar yield of all cell and EV preparations (Sup. Fig 7A). As expected, decreasing concentrations of hTERT-plasmid input DNA resulted in higher ct values and water control gave no amplification signal after 35 amplification cycles (Sup. Fig 7B). We then confirmed the presence of hTERT plasmid DNA in genomic DNA preparations of WJ-MSC/TERT273 showing a slightly, but significantly higher hTERT DNA content compared to their primary counterparts. Signals were close to qPCR detection limits (Sup. Fig 7C). No hTERT plasmid DNA was observed in TERT EVs when compared to primary cell derived EVs (Fig 6B).

To further confirm that hTERT immortalization of cells does not result in conferring telomerase activity to the EVs, we performed RT-TRAP assays to measure relative telomerase activity (TA) of cells and EVs normalized to HEK293 cells (Fig. 6C). While WJ-MSC/TERT273 cells show a robust telomerase activity, neither primary nor telomerized MSC-EVs showed detectable TA.

Finally, we tested if ectopic expression of hTERT would allow telomerized WJ-MSCs cells to grow in an anchorage-independent way using soft agar colony formation assay as an in vitro tumorigenicity assay. 6 days after seeding 3000 and 9000 cells/well in soft agar, no cell proliferation or colony formation was visible (Fig. 6D) or measurable (Fig. 6E, 6F) upon CyQuant GR Dye based colony staining for primary and telomerized WJ-MSCs, even at late population doublings, as compared to HEK293 cells used as positive control. The absence of acquired anchorage-independent cell growth or TA activity suggests that ectopic expression of hTERT does not change the safety profile of telomerized cells as compared to their primary counterparts.

## DISCUSSION

In this study, we aimed to explore the impact of hTERT-mediated immortalization (‘telomerization’) of MSCs on the characteristics of their secreted EVs, given the increasing demand for human cell factories to produce complex biopharmaceuticals (Fliedl et al., 2015) due to the advent of EV-based technologies.

Immortalization of producer cells while maintaining them as similar as possible to primary cells seems a highly promising strategy to overcome some of the limitations of using primary MSCs as cell sources for EV manufacturing, such as limited lifespan, heterogeneity, batch-to-batch or donor-to-donor variability (Gimona et al., 2021; Johnson et al., 2021). Of these, elimination of donor heterogeneity of producer cells seems one of the most important steps to advance scalable and standardized EV production for therapeutic applications (Labusek et al., 2023; Roura and Bayes-Genis, 2019; Witwer et al., 2013).

One possibility for generation of high numbers of homogenous MSCs is to differentiate them from induced pluripotent stem cells. Such iMSCs are considered young and can support e.g. osteogenic differentiation by their ECM and secretome (Hanetseder et al., 2023). However, they also were reported to display some functional differences compared to tissue-derived MSCs, such as lower immunomodulatory potential (Bertolino et al., 2022; Frobel et al., 2014) or hampered adipogenic differentiation capacity (Chen et al., 2012; Diederichs and Tuan, 2014). Therefore, we here used hTERT as an immortalization agent. We first confirmed that the telomerization of WJ-MSC273 enabled the cells to bypass senescence. Then, we demonstrated that during the theoretically unlimited in vitro propagation of the WJ-MSC/TERT273, cells maintained a constant doubling time and a homogeneous cell morphology over passages, resembling early PD WJ-MSC273 primary cells. During replicative ageing, the MSC phenotype changes not only in terms of morphology and proliferation, but also in terms of functionality (Adlerz et al., 2020; Izadpanah et al., 2008; Kretlow et al., 2008; Lo Surdo and Bauer, 2012) 201. Hence, we tested if telomerization of MSCs would affect either the cell characteristic surface antigen profile or the trilineage differentiation potential. WJ-MSC/TERT273 did not show changes in differentiation potential over time, while primary WJ-MSCs underwent growth arrest in accordance with previous reports on senescence of MSCs (Wolbank et al., 2009). Thus, both WJ-MSC273 and telomerized cells fulfill the minimal criteria of human MSCs, showing sufficient adherence to plastic during cell culture, expression of CD105, CD73 and CD90, while lacking CD34 and showing sustained differentiation potential towards osteoblasts, adipocytes and chondroblasts in vitro as defined by the International Society for Cell & Gene Therapies (Dominici et al., 2006; ISCT.; Viswanathan et al., 2019). This is not achieved when induced pluripotent stem cells (iPSCs) are used for the large-scale generation of standardized MSCs (Bertolino et al., 2022).

Moreover, not only the cells, but also the cargo and biological activity of MSC-EVs changes with increased replicative age of the cells (Boulestreau et al., 2020). For example, late PD MSC-derived EVs showed a decreased vasculogenic activity (Patel et al., 2017), a reduction of the osteogenic activity (Miclau et al., 2023) and an increase in paracrine senescence and apoptosis of treated cells as compared to early PD MSC-derived EVs (Khayrullin et al., 2019). It has also been demonstrated that EVs from late passage MSCs show lower immunomodulatory potential in a mouse model of ocular Sjögren’s syndrome (Kim et al., 2020), as well as a reduced protective effect in LPS-induced acute lung injury in young mice (Huang et al., 2019). To overcome these issues, different strategies have been exploited to immortalize the parental MSCs for EV production (Chen et al., 2011; Hindle et al., 2024; Labusek et al., 2023). Although the resulting EVs induced cell proliferation and were cardio- (Chen et al., 2011) and neuroprotective (Labusek et al., 2023), some immortalization strategies using oncogene overexpression resulted in changes in the cellular phenotype of EV producing cells (Chen et al., 2011). It is important to note that the loss of MSC characteristics in MYC-immortalized producer cells may influence the properties of secreted EVs and may lead to a transfer of oncogenes to target cells, which must be carefully monitored. On the other hand, it has been described that large-scale production of EVs from a previously established, telomerized adipose derived MSC line shows very limited batch-to-batch variability between different EV preparations, as well as similar reparative properties against damaged retinal pigment epithelial cells to those observed in primary MSC EVs (Hindle et al., 2024).

Considering that the field still lacks surrogate markers to predict the inherent therapeutic potency of MSCs or of its secreted particle preparations (Reiner et al., 2017; Witwer et al., 2019) a comprehensive characterization of the primary and telomerized WJ-MSC-derived EVs was necessary. No differences in particle release, particle size, total protein content or surface marker profile were noticeable between primary and immortalized WJ-MSC EVs. Similarly, we demonstrated that the miRNA cargo of EVs from WJ-MSC273 and WJ-MSC/TERT273 cells was highly similar. Interestingly, the correlation analysis revealed an upregulation of hsa-miR-148a-3p and has-miR-335-3p in telomerized WJ-MSC derived EVs; both miRNAs are involved in the negative regulation of adipogenic differentiation of MSCs, cholesterol homeostasis and clearance and bone remodeling (Lincoln et al., 2022; Manochantr et al., 2017; Tomé et al., 2014). On the other hand, both miRNAs are involved in changes of the cytoskeletal organization (Uray et al., 2020). In this regard, it has been proven that exosomal miR-148a-3p in EVs from human placental mesenchymal stromal cells can attenuate endothelial barrier dysfunction by promoting cytoskeletal remodeling (Lv et al., 2024), while exosomal miR-335-3p from bone marrow mesenchymal stromal cell promoted bone fracture recovery and osteoblast differentiation by releasing miR-335; this effect was impaired by miR-335 downregulation in BM-EVs (Hu et al., 2021). Overall, despite the fact that paracrine activities of MSC-EVs due to their miRNA cargo is disputed (Albanese et al., 2021; Qiu et al., 2018), because of low abundance of miRNAs in EVs (Chen et al., 2016; Diendorfer et al., 2022), these results suggest that the miRNA content of telomerized EVs is very similar to the primary counterpart, and might confer similar, if not improved biological activities.

To validate the EV activity, functional biological assays were performed to assess the inherent anti-inflammatory, anti-fibrotic and tissue regenerative properties typical of MSC-EVs. The results revealed no significant differences in biological activities, with similar dose responses observed. The particle concentrations tested in these assays had previously been reported in other functional MSC-EV studies (Pacienza et al., 2019), showing comparable anti-inflammatory properties. These findings are noteworthy, as changes in other parameters, such as culture conditions, appear to have a more pronounced impact on EV potency compared to cellular immortalization via hTERT (Shekari et al., 2023).

Additionally, 3D culturing systems have been shown to significantly influence the metabolome of EVs (Palviainen et al., 2019) and alter MSC-EV cargo composition, potentially hampering their biological functions (Gobin et al., 2021; Keysberg et al., 2023; Kusuma et al., 2022; Rocha et al., 2018; Yan and Wu, 2020). Hence, as telomerized MSCs seem to exhibit a behavior similar to their primary counterparts, culture conditions that enhance EV production might be easily adaptable to them. For example, serum-free media (Almeria et al., 2022; Haraszti et al., 2019; Li et al., 2015), but also xeno-free media (Scheiber et al., 2022) have been shown to boost EV production per cell. MSC-EVs from cells cultured in hypoxic conditions show significant changes in cargo composition and increased angiogenic properties compared to EVs derived from normoxic MSCs (Ge et al., 2021; Gregorius et al., 2021; Zhang et al., 2012)

Finally, we aimed to test key safety aspects of WJ-MSC/TERT273, and the secreted particles. It has been well recognized that human telomerase is not an oncogene and not sufficient to drive cell transformation, since additional chromosomal aberrations and oncogenic transformation in telomerized cells are needed to form tumorigenic cells (Burns et al., 2017; Elenbaas et al., 2001). Similarly, whole mouse body hTERT overexpression has been proven to be therapeutically beneficial in anaemic mice (Bär et al., 2016), aged mice (Muñoz-Lorente et al., 2018), a mouse model of myocardial infarction (Bär et al., 2014) and in one of pulmonary fibrosis (Povedano et al., 2018) with no increased tumor formation reported. Additionally, numerous studies have shown that hTERT immortalized cells (Chang et al., 2005; Iacomi et al., 2022; Liu et al., 2013; Simonsen et al., 2002; Wieser et al., 2008), as well as MYC transformed MSCs, do not induce anchorage-independent cell growth in vitro nor form tumors when injected subcutaneously in vivo (Tan et al., 2021). Taking these reports into consideration, even if trace amounts of hTERT DNA or mRNA would be present in EVs derived from telomerized cells, no biosafety concern is expected. Still, when analyzed, no enrichment of small RNAs mapping onto human telomerase was observed in primary and TERT EVs, nor was telomerase activity detected. In line with the absence of acquired anchorage-independent growth of primary or telomerized cells, our data suggest that immortalized MSCs can be regarded as safe cell factories producing bioactive EVs. Similar data were observed for MYC-immortalized MSCs by Sai Kiang Lim’s group (Tan et al., 2021), proving that not only immortalised cells, but also the secreted EVs do not show tumorigenic potential. Daily intraperitoneal injections of sEVs from MYC immortalized MSCs into athymic nude mice with human head and neck cancer xenografts for 28 days did not promote tumor growth.

Overall, these results suggest that non-viral immortalization of MSCs by hTERT overexpression does not affect the inherent molecular, biological, and functional properties of either donor cells or MSC-derived EVs tested here. Thus, we suggest the use of telomerized MSCs instead of primary cells as EV cell factories promoting scalable EV production, while enabling higher standardization of the generated EV product by decreasing donor-to-donor and even lot-to-lot variability (Chance et al., 2019; Kodama et al., 2022; Kordelas et al., 2019; McBride et al., 2021; Ragni et al., 2019; Scheiber et al., 2022).

In addition, stable cell lines offer various advantages over primary cells. It has been demonstrated that the immortalization of a mixed, polyclonal population of primary cells could lead to the generation of heterogeneous cell lines (Carius et al., 2023). However, a monoclonal selection strategy might improve the therapeutic and functional homogeneity and properties of the immortalized cell line. In addition, telomerized cells can be engineered, as we recently demonstrated for our kidney epithelial RPTEC/TERT1 cell line. The cells were first gene edited and then subcloned to a monoclonal cell line, while the primary cell-like characteristics were maintained (Wieser et al., 2019), a sequence of methods hardly feasible using primary cells. Thus, this strategy opens a wide range of endogenous genetic engineering possibilities to alter tropism (Vogt et al., 2021), therapeutic cargo of EVs or the use of engineered EVs including e.g. tags for affinity purification and tracking (Bobbili et al., 2024; Monroe et al., 2024). All of this is conferring specific additive functions to EVs for advancing their inherent therapeutic capabilities.

Taken together, we here suggest that telomerized MSCs are fit for purpose as human cell factories with a huge potential to customize the resulting EVs and produce them in scalable, standardized ways as a basis for clinical translation.

## ACKNOWLEDGEMENTS

We thank Kitti Dora Csalyi at the Max Perutz Labs BioOptic-FACS Core Facility for the support with flow cytometric analysis. We thank Gabriel Brognara at the Protein Technologies Facility (ProTech) of Vienna BioCenter Core Facilities for the support with Western Blot analysis. In addition, this project was supported by the BOKU Core Facility for Biomolecular and Cellular Analysis This study was supported by the European Union within the Horizon Europe MSCA program Cellular Homeostasis ANd AGing in Connective TissuE Disorders (CHANGE, grant agreement No 101072766).

## AUTHOR CONTRIBUTIONS

Alessia Brancolini: Conceptualization; data curation; formal analysis; investigation; methodology; visualization; validation; writing—original draft; writing—review and editing. Madhusudhan Reddy Bobbili: Conceptualization; formal analysis; investigation; methodology; validation; supervision. Marianne Pultar: Formal analysis; data curation; software; writing—review and editing. Zahra Mazidi: Investigation; methodology; validation. Matthias Wieser: Investigation; methodology; validation. Johanna Gamauf: Investigation; methodology; validation. Marieke Roefs: Methodology; validation. Giulia Corso: Methodology; validation. Harini Nivarthi: Methodology; validation. Maria Belen Arteaga Paredes: Investigation; methodology. Dieter Bettelheim: Resources. Elsa Arcalis: Investigation; methodology. Sivun Dmitry: Investigation; methodology. Jaroslaw Jacak: Conceptualization; supervision. Iris Gerner: Conceptualization; supervision. Matthias Hackl: Conceptualization; supervision. Johannes Grillari: Conceptualization; project administration; supervision; writing—original draft; writing—review and editing. Regina Grillari-Voglauer: Conceptualization; project administration; supervision; writing—original draft; writing—review and editing. All authors have read and approved the manuscript.

## DATA AVAILABILITY STATEMENT

The paper and its supplementary materials provide the data that support the findings of this study.

## SUPPLEMENTARY FIGURES

**Supplementary Figure 1.**
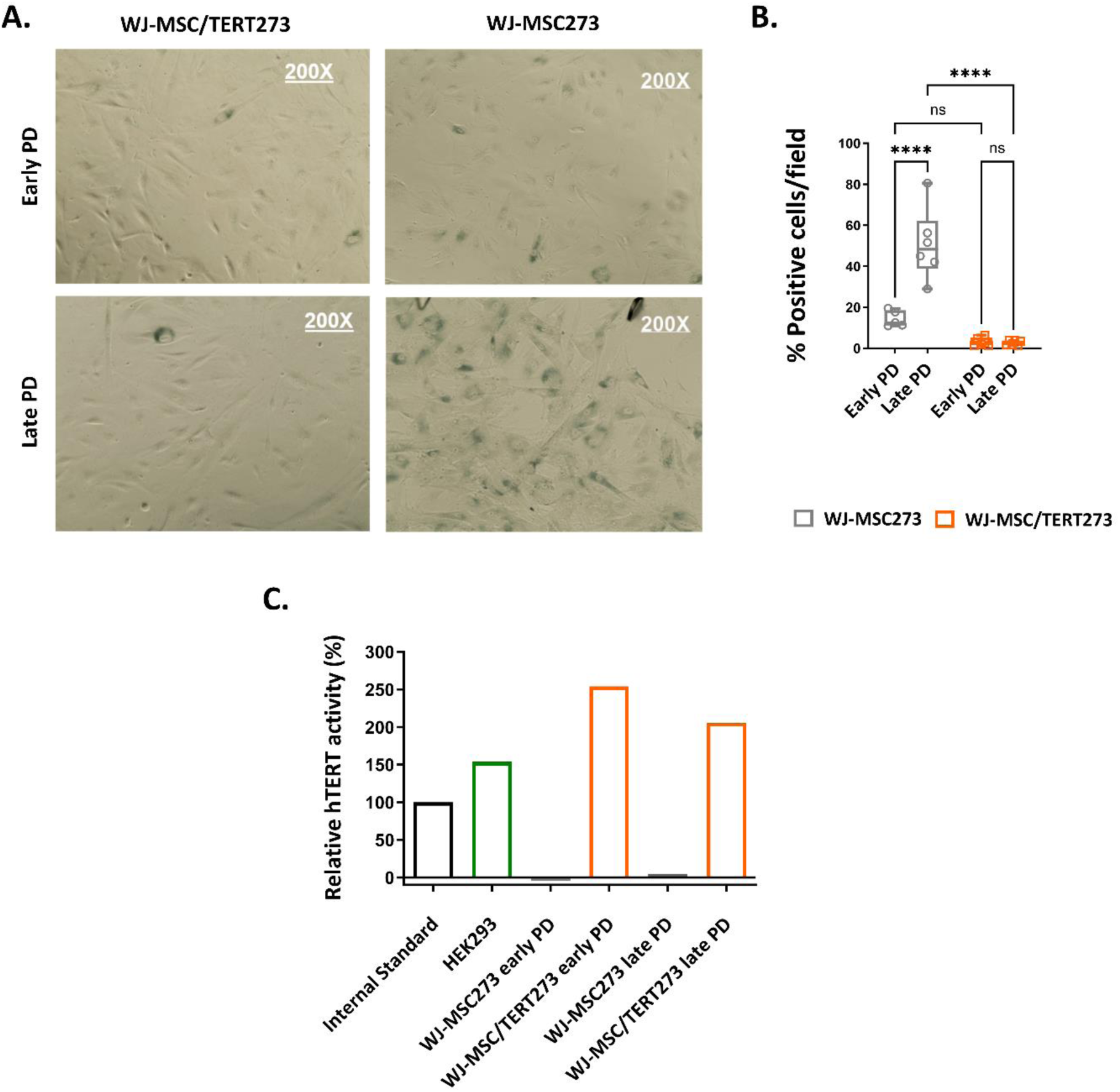
**A**. Representative microscopy images and **B.** quantification of SA-β-Galactosidase staining of primary and telomerized WJ-MSCs at early (PD23.4 and 28.5) and late population doubling times (PD64,2 and 60.6) by bright field microscopy. Images were taken at 200X magnification. Significance levels are reported as follows: **** p < 0.0001. The data are presented as mean ± SD (n=3; 2 photos/well). **C.** Evaluation of telomerase activity in primary and telomerized WJ-MSCs at early and late PDs by TeloTAGGG Telomerase PCR ELISAPLUS assay and TA: Primary cells were tested at PD 11.0 and 54.2, while WJ-MSC/TERT273 were tested at PD18.6 and 58.4; the relative hTERT activity was determined in comparison to the internal Control Template TSR8. Telomerase activity in HEK273 cells was measured as positive control.

**Supplementary Figure 2.**
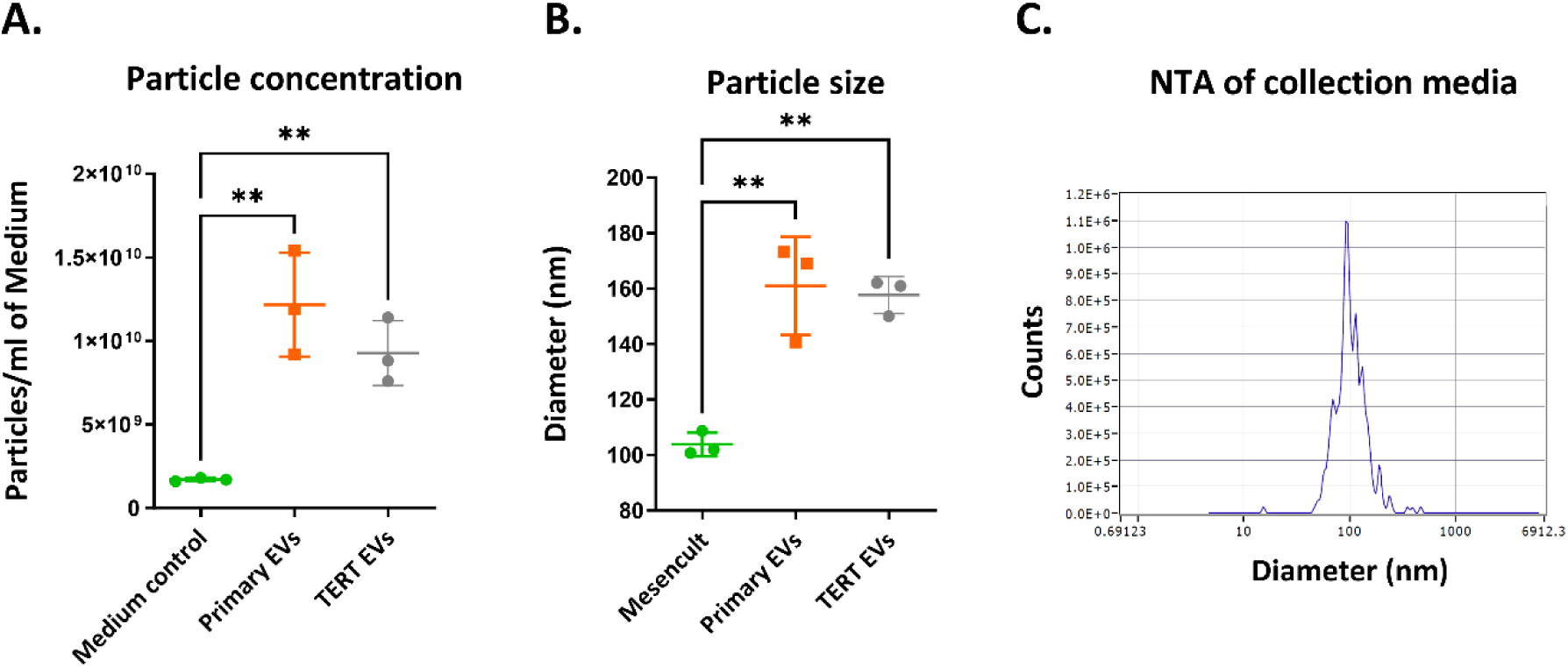
**A.** Particle concentration from EV-collecting medium (Mesencult ACF, left), primary (center) and telomerized (right) WJ-MSC derived EVs. For all conditions, 45 mL of EV-collecting medium or conditioned medium were enriched by TFF (45:1 enrichment ratio) and measured by NTA. **B.** Mean particle diameter from EV-collecting medium (Mesencult ACF, left), primary (center) and telomerized (right) WJ-MSC-derived EVs by NTA. For particle concentration and particle size, significance levels are reported as follows: ** 0.001 ≤ p < 0.01; The data are presented as mean ± SD (n=3). **C.** Representative size distributions of particles from EV-collecting medium (Mesencult ACF, left) by NTA.

**Supplementary Figure 3.**
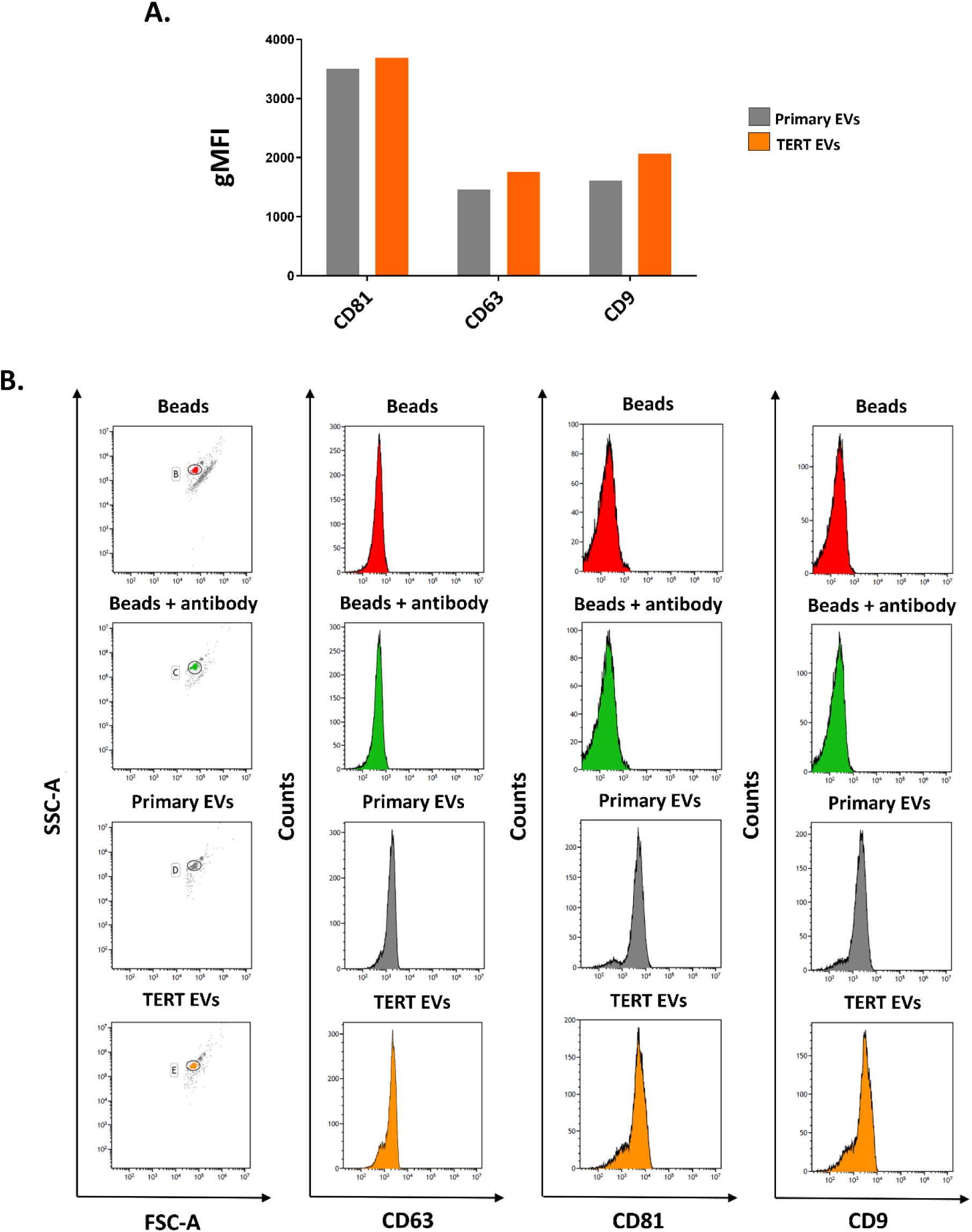
**A.** Quantification and comparison of the presence of canonical EV-enriched tetraspanins and **B.** assessment of canonical EVs-enriched tetraspanin expression on primary and telomerized WJ-MSCs-derived EVs by CD81+ beads-based flow cytometry analysis.

**Supplementary Figure 4.**
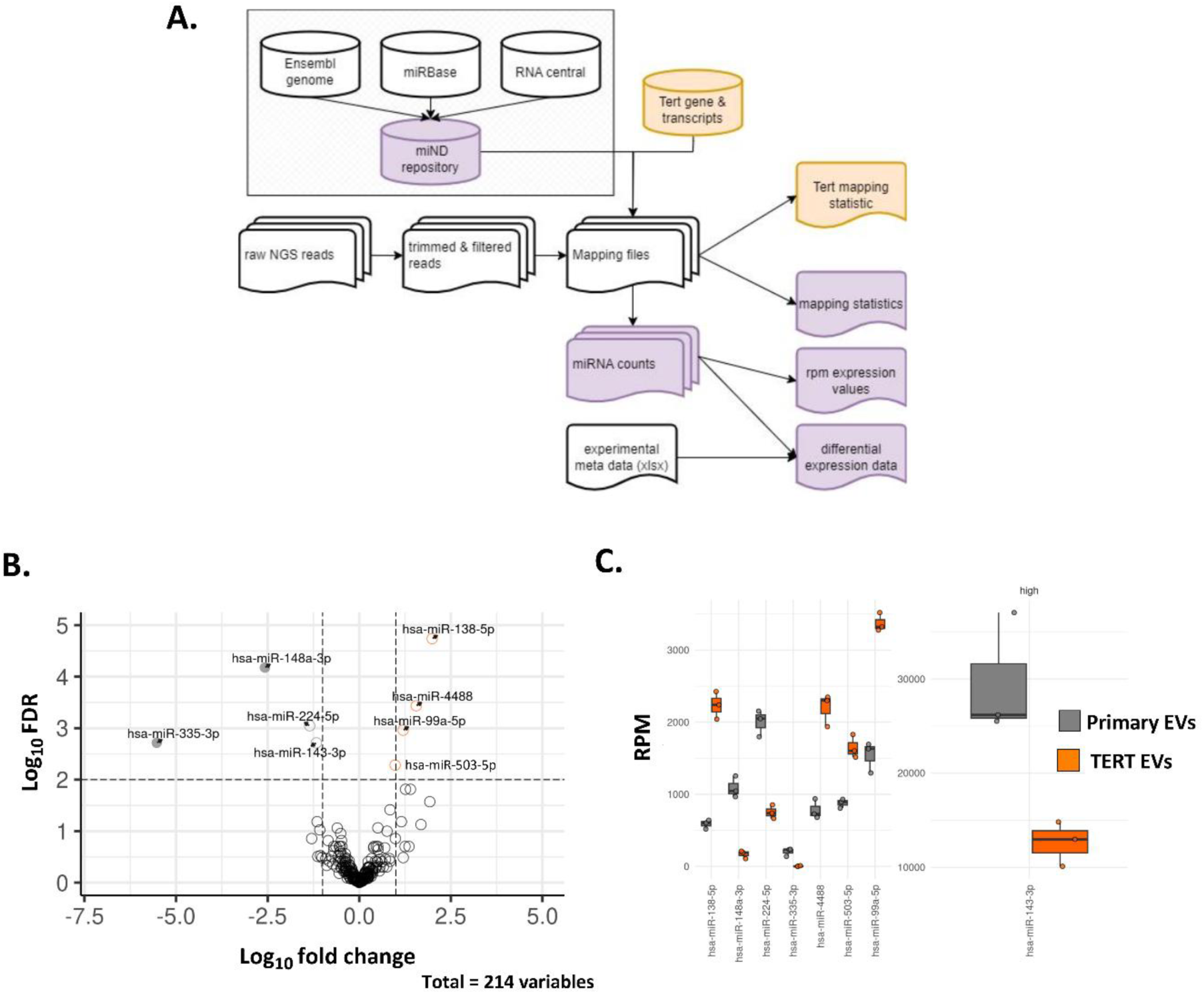
**A.** Experimental workflow of the small RNA sequencing and analysis of primary and TERT EVs. **B.** Differentially expressed miRNAs between primary and hTERT-MSC EV-cargo. Volcano Plot displaying enriched miRNAs with a FDR < 0.01 in TERT EVs (log2 fold change > 1, orange) and primary EVs (log2 fold change < -1, grey, log2 fold change < -2, filled grey). **C.** RPM expression of the 8 miRNAs found to be differentially expressed in primary and TERT EVs (log2 fold change < -1’ log2 fold change > 1 and FDR < 0.01).

**Supplementary Figure 5.**
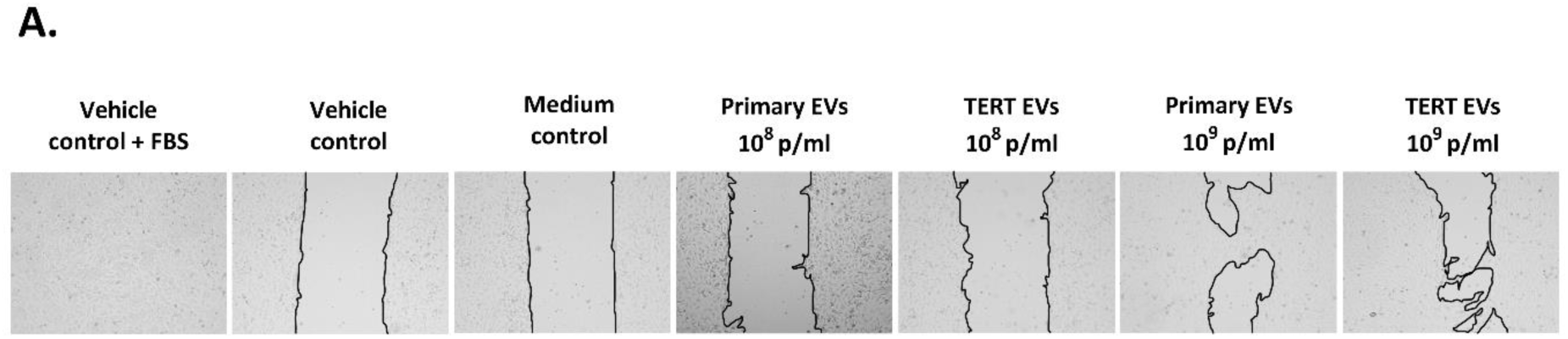
**A**. Representative microscopy images of the wound closure of HUVEC/TERT2 cells 40 h after the removal of culture insert and treatment.20 mM HEPES (vehicle control) and TFF enriched cell-free EV collection medium (medium control) were included as negative controls, while complete culture medium (vehicle control + FBS) was included as positive control

**Supplementary Figure 6.**
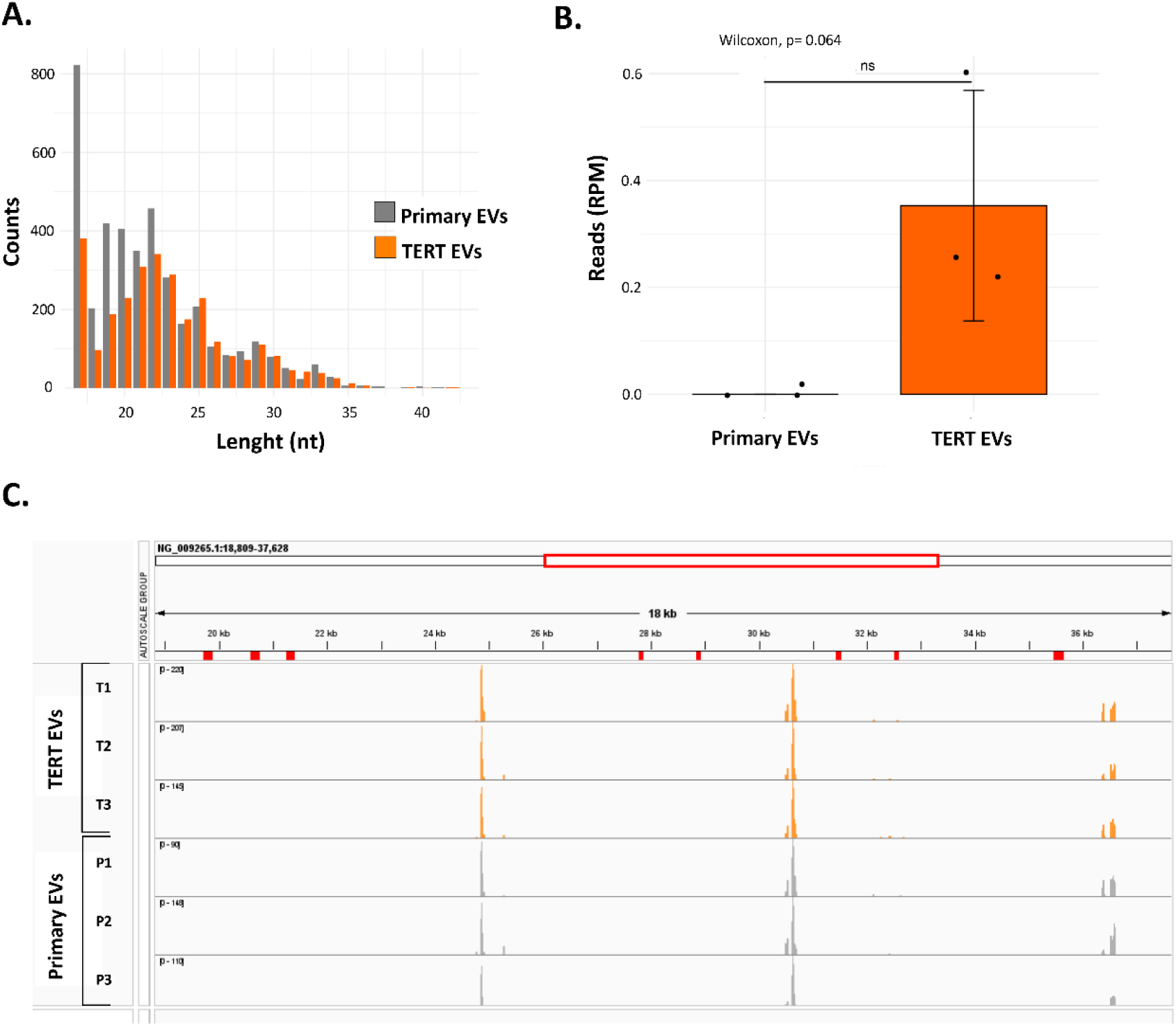
**A**. Length distribution of reads mapping against the gene sequence of human telomerase (NG_009265.1, NM_001193376.3 and NM_198253.3) in primary and TERT EVs. **B.** Normalized number of mapped reads (RPM) against the human telomerase transcript sequence (NM_198253.3). **C.** Representative snapshot of reads mapping to the gene sequence of human telomerase (NG_009265.1) in primary (P1, P2, P3) and TERT EVs (T1, T2, T3). Exonic regions are highlighted in red.

**Supplementary Figure 7.**
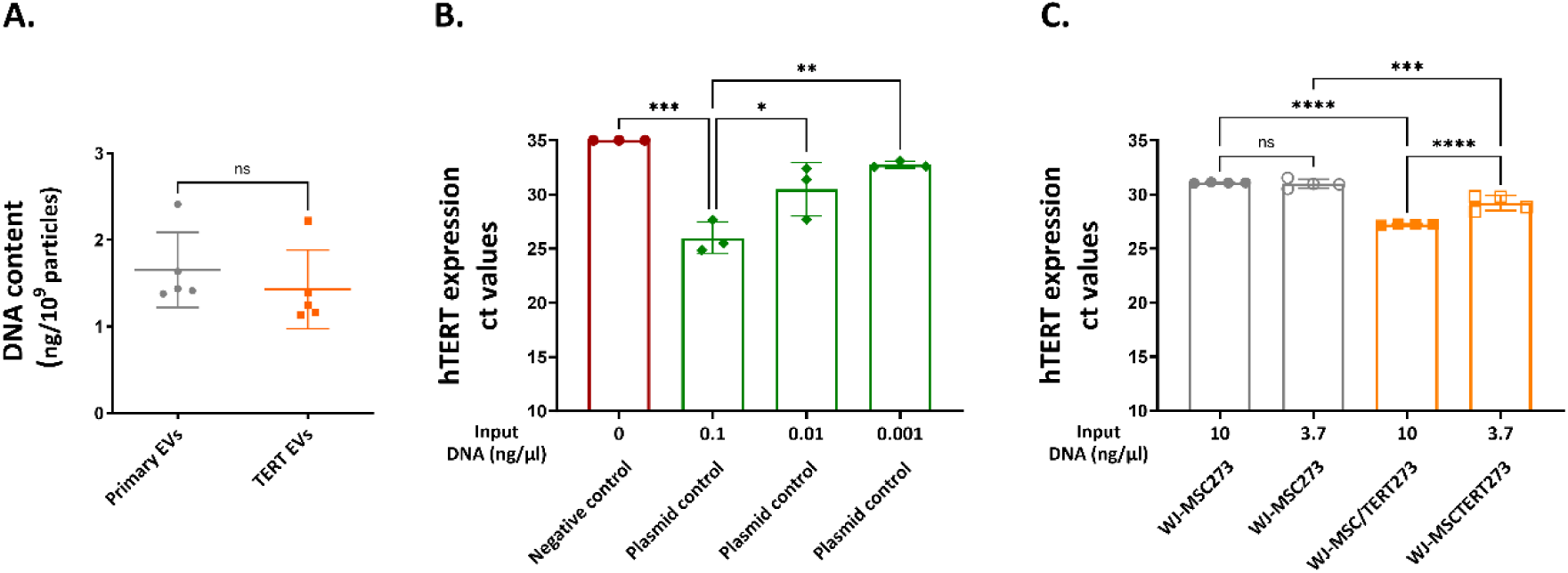
**A.** Quantitation of DNA extraction from WJ-MSC273 and WJ-MSC/TERT273 derived EVs. DNA extraction was performed using Qiamp DNA Micro kit and the concentration of extracted DNA was normalized on final elution volume and particle concentration (estimated by NTA) of the original EV-containing samples. **B.** Evaluation of hTERT qPCR results on hTERT-carrying plasmid DNA. The graph shows the individual ct-values from hTERT-carrying plasmid DNA tested at input concentrations of 0.1 ng/µl, 0.01 and 0.001 ng/µl. For negative control (water control), lack of amplification by qPCR after 35 cycles was represented as ct value = 35 **C.** Evaluation of hTERT qPCR results on genomic DNA from primary and telomerized WJ-MSCs. The graph shows the individual ct-values from WJ-MSC273 and WJ-MSC/TERT273 DNA tested at input concentrations of 10 ng/µl and 3.7 ng/µl. A negative cut-off value was set at ct value = 35. Significance levels are reported as follows: * 0.01 ≤ p < 0.05; ** 0.001 *** 0.001 ≤ p < 0.0001; **** p < 0.0001. The data are presented as mean ± SD, with n=5 for DNA extraction efficiency, as mean ± SD, with n=4 for qPCR results on hTERT genomic DNA and n=3 for qPCR results on -hTERT-carrying plasmid DNA.

